# Satisfying leaf nitrogen demand under drought is determined by soil nitrogen availability and drought severity

**DOI:** 10.1101/2025.03.11.642652

**Authors:** Alissar Cheaib, Nicholas G. Smith

**Affiliations:** Department of Biological Sciences, Texas Tech University, Lubbock, TX, USA

**Author notes:** Correspondence to: Experimental Sciences Building 2, Texas Tech University 1006 Canton Ave, Lubbock, TX 79409 USA. **Author contributions** AC and NGS conceived the study. AC conducted the experiment, AC and NGS performed the data synthesis and drafted the manuscript. **Data statement** All data and code used for these analyses are available at: https://doi.org/10.5281/zenodo.15000895.

**Keywords:** Photosynthetic least-cost theory, soil drought, leaf nitrogen, leaf water-use efficiency, carbon costs of resource acquisition, V_cmax_, A_net_

## Abstract

Plants balance water and nitrogen acquisition to optimize carbon gain, particularly under resource-limited conditions. These tradeoffs can be predicted from the photosynthetic least-cost theory, but its applicability under severe drought and nitrogen limitation within a single species remains untested.

We examined how soil water stress and nitrogen limitation shape leaf-level water and nitrogen economics and the associated plant carbon costs of soil resource acquisition in common sunflower (*Helianthus annuus* L.) grown under two nitrogen levels and three soil water availability scenarios.

Contrary to photosynthetic least-cost theory, increasing drought severity reduced leaf nitrogen content (N_area_) under low nitrogen, likely due to impaired nitrogen uptake via transpiration-driven mass flow. Likewise, the maximum Ribulose-1,5-Biphosphate Carboxylase/Oxygenase (RuBisCO) carboxylation rate (V_cmax_) and net photosynthetic rate (A_net_) decreased over time as drought progressed under low soil nitrogen. However, under high nitrogen, and in line with least-cost theory, N_area_, V_cmax_ and A_net_ remained stable, with N_area_ increasing over time, enhancing leaf water-use efficiency.

Total leaf area and biomass increased with soil nitrogen availability, but this effect weakened under severe drought. Notably, carbon costs of nitrogen acquisition increased with drought severity only under low nitrogen, indicating water stress impedes nitrogen uptake. These findings highlight the need to integrate soil resource demand and availability into photosynthesis models, especially under extreme drought and nitrogen limitation.

## Introduction

Throughout their evolutionary history and lifetimes, land plants, as sessile organisms, have faced complex and interacting abiotic and biotic stresses (Du *et al*., 2024). These multiple stressors, often acting in concert, have shaped plant functional traits, reflecting trade-offs and strategies for resource acquisition that have determined both acclimatory adjustments and long-term evolutionary trajectories (Westoby *et al*., 2002; Edwards *et al*., 2010; Reich, 2014; Volaire, 2018; VanWallendael *et al*., 2019; Volaire *et al*., 2020).

One prominent example of these trade-offs is the balance between water loss and carbon gain. Under drought conditions, plants partially close their stomata to conserve water (Buckley, 2005; Buckley *et al*., 2017). Most terrestrial plants have evolved tight stomatal control over water loss, preserving the integrity of the soil-plant hydraulic continuum (Wankmüller & Carminati, 2023; Torres Ruiz *et al*., 2024). However, while this partial stomatal closure reduces transpiration, it also reduces CO_2_ influx, lowering the intracellular CO_2_ concentration (C_i_) and net photosynthetic rate (Brodribb, 1996; Flexas *et al*., 2014; Lawson & Vialet Chabrand, 2019). Since evolution has not produced a membrane permeable to CO_2_ but impermeable to water (Cowan, 1978), land plants face an inherent “dilemma” (Raschke, 1976) forcing them to adjust trade-offs between water conservation and carbon assimilation.

To optimize water-use efficiency, plants adjust their rate of photosynthesis relative to transpiration, balancing carbon gain with the costs of maintaining photosynthetic capacity and water transport capacities (Prentice *et al*., 2014; Wang *et al*., 2017). But how do plants achieve this optimization? According to photosynthetic least-cost theory (Wright *et al*., 2003; Prentice *et al*., 2014; Franklin *et al*., 2020; Harrison *et al*., 2021), water and nitrogen (N) are substitutable: a given photosynthetic rate can be achieved through different combinations of leaf nitrogen content per area (N_area_) and stomatal conductance (g_s_). Nitrogen plays a central role in this trade-off because it is critical for photosynthesis, with more than 70% of leaf nitrogen allocated to photosynthetic enzymes such as Ribulose-1,5-Biphosphate Carboxylase/Oxygenase (RuBisCO) (Evans, 1989; Evans & Seemann, 1989; Onoda *et al*., 2017; Evans & Clarke, 2019). Under drought conditions, reduced g_s_ requires a greater investment in nitrogen acquisition and allocation to the photosynthetic machinery in leaves to enhance the maximum rate of RuBisCO carboxylation (V_cmax_) and CO_2_ fixation, compensating for the reduced intercellular CO_2_ concentration. In contrast, wet conditions favor a lower investment in leaf nitrogen and V_cmax_.

Consequently, the regulatory role of nitrogen introduces a trade-off between leaf water-use efficiency (WUE_leaf_) and leaf nitrogen-use efficiency (NUE_leaf_) (Patterson *et al*., 1997; Zhong *et al*., 2019). A decrease in g_s_ and water loss, coupled with increased investment in leaf nitrogen under drought, enhances WUE_leaf_ at the expense of NUE_leaf_, whereas the opposite trade-off occurs under wet conditions. Stomatal conductance reflects the cost of water supply for transpiration, while leaf nitrogen content represents the investment in V_cmax._ The unit costs of nitrogen and water acquisition determine the ratio of intracellular to ambient CO_2_ concentration (C_i_/C_a_, or χ). A low χ value indicates either a relatively low g_s_, an increased photosynthetic capacity that reduces C_i_, or a combination of both (Condon, 2004).

Following the photosynthetic least-cost theory, the unit costs of nitrogen and water acquisition are captured by a ratio (β_leaf_) (Prentice *et al*., 2014; Wang *et al*., 2017; Smith *et al*., 2019; Smith & Keenan, 2020; Stocker *et al*., 2020; Paillassa *et al*., 2020):

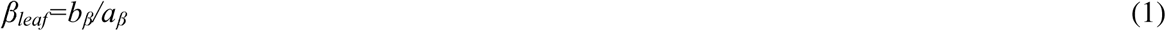

where *b*_β_ represents the carbon cost of acquiring nutrients and maintaining photosynthetic enzymes to support carboxylation and *a*_β_ represents the carbon cost of acquiring water and maintaining water transport to support assimilation at the same rate. The term *b*_β_ is expected to increase with decreasing nutrient availability, as acquiring nutrients in low-resource conditions becomes more costly. In contrast, the term *a*_β_ is expected to increase with increasing drought severity, because the cost of acquiring water becomes higher when it is scarce compared to when it is abundant. This ratio (β_leaf_) is considered at the leaf level and assumed to scale up to the whole plant, although the validity of this assumption has not been well tested (Stocker *et al*., 2025).

Many observational and experimental studies have shown that dryland species often exhibit higher leaf nitrogen content and V_cmax_ than species from wetter ecosystems, supporting the least-cost hypothesis and underscoring the consistent response of plants in adjusting their leaf nitrogen content to aridity (Domingues *et al*., 2010; Prentice *et al*., 2014; Ferreira Domingues *et al*., 2015; Smith *et al*., 2019; Peng *et al*., 2021; Luo *et al*., 2021; Querejeta *et al*., 2022; Westerband *et al*., 2023; Fan *et al*., 2024; Cheaib *et al*., 2025). However, since the nitrogen uptake by roots is tightly coupled with soil moisture (Leadley *et al*., 1997; McMurtrie & Näsholm, 2018), this raises a key question: how do plants enhance leaf nitrogen content to counteract reduced g_s_ when the soil dries?

As plants rapidly deplete nutrients from the rhizosphere, replenishment depends on dissolved nutrients being transported via the transpiration-driven mass flow of water to the root zone (Voltas *et al*., 1998; Cabrera Bosquet *et al*., 2009; Lambers *et al*., 2014). However, a reduction in g_s_ diminishes the transpiration stream, restricting the movement of water and low-molecular-weight solutes (e.g., nutrient ions, organic acids). Consequently, the transport and diffusion of solutes from the bulk soil to root surfaces via mass flow become limited (Barber, 1966; Kreuzwieser & Gessler, 2010; Chapman *et al*., 2012; Lambers & Oliveira, 2019). This dependence on mass flow for soil nutrient transport is widely recognized and forms a core component of models simulating soil nutrient fluxes (Nye & Marriott, 1969; Rengel, 1993). Many studies have highlighted the importance of transpiration-driven mass flow in delivering nitrogen to the root surface (Barber, 1962; Tanguilig *et al*., 1987; Kirnak *et al*., 2002; Cramer *et al*., 2009; Waraich *et al*., 2011; Lambers *et al*., 2014; Sardans & Peñuelas, 2014), particularly nitrate—the primary form of soluble inorganic nitrogen absorbed by roots. Some studies have even reported that when plants are unable to access nitrogen through interception or diffusion, they increase their stomatal conductance and transpiration as a strategy to acquire nitrogen via mass flow (Cramer *et al*., 2008; Matimati *et al*., 2014). Drought has been reported to reduce the abundance and activity of major uptake proteins for nitrogen and phosphorus, explaining the lower nutrient uptake rates by roots (Bista *et al*., 2018, 2020). A meta-analysis assessing the impact of drought on plant nutrients (He & Dijkstra, 2014) found that soil drought significantly reduced total plant nitrogen and phosphorus in short-term studies (<90 days), especially under prolonged, uninterrupted drought stress. The authors attributed this decline in plant nutrients to the direct impact of drought on nutrient uptake rather than a reduction in nutrient availability. If transpiration-driven mass flow plays a key role in nitrogen acquisition, can photosynthetic least-cost theory be generalized across different types and levels of soil water stress and soil nitrogen availability? Moreover, how would a reduction in transpiration-driven mass flow influence the carbon cost of nutrient acquisition and water acquisition? Understanding how these two costs interact is crucial, as their interplay could directly shape the nitrogen-water tradeoffs in leaves, influencing photosynthetic capacity through the balance between χ, N_area_ and V_cmax_. To our knowledge, no study has explored these questions across varying levels of soil water stress and soil nitrogen availability.

To explore these questions, we grew common sunflower (*Helianthus annuus* L.) plants in large, 18 L pots for approximately 70 days. Plants were subjected to a full factorial combinations of nitrogen availability (two levels) and soil water availability (three levels). The three scenarios included well-watered, slow moderate progressive drought, and fast severe developing drought with one re-wetting cycle (see Materials and Methods section). We quantified the functional leaf and photosynthetic traits throughout drought progression at four time points, along with the biomass and the costs of nutrient and water acquisition at whole-plant scale.

We chose these drought scenarios because we expected that the severity, frequency, and duration of soil drought would interact with soil nitrogen levels to affect a plant’s ability to meet its leaf nitrogen demand and the relative costs of acquiring nitrogen and water. Based on these considerations, we proposed the following hypotheses:

i. **Hypothesis 1**: As drought progresses throughout the experiment, we hypothesized that N_area_ in severely drought-stressed plants, particularly under the fast-developing drought scenario, would increase compared to well-watered plants, in accordance with photosynthetic least-cost theory, but only for plants grown in soils with high nitrogen availability (Figure 1a). In contrast, and because the rate of solute transport is influenced by solute concentration (Nielsen *et al*., 1986), we hypothesized that plants subjected to low nitrogen availability would experience limitations in nitrogen uptake, insufficient to meet leaf nitrogen demand or compensate for the effects of partial stomatal closure. As a result, N_area_ in drought-stressed plants with low nitrogen availability was expected to decrease over the course of the experiment compared to well-watered plants (Figure 1a), preventing photosynthetic least-cost theory from holding.
ii. **Hypothesis 2**: In accordance with photosynthetic least-cost theory, we expected plants grown under high nitrogen availability to show an increase in water-use efficiency (WUE_leaf_) and a decrease in nitrogen-use efficiency (NUE_leaf_) with increasing drought severity (Figure 1b). However, because plants grown under low nitrogen availability were expected to be unable to meet their leaf nitrogen demand as soil drought severity increases, we did not anticipate an increase in WUE_leaf_ with increasing drought severity (Figure 1b), contrary to photosynthetic least-cost theory. Consequently, water-stressed plants grown under high soil nitrogen levels were expected to exhibit higher WUE_leaf_ compared to those grown under low soil nitrogen levels.
iii. **Hypothesis 3**: At the whole-plant level, plants grown under low soil nitrogen availability were expected to incur a higher carbon cost for nitrogen acquisition (N_cost_) compared to those in high nitrogen levels, as nitrogen becomes increasingly scarce and costly to obtain at low soil concentrations (Figure 2, arrow a). Additionally, if drought directly hinders nitrogen uptake by reducing mass flow, we expected drought severity to further increase the carbon cost of nitrogen acquisition (Figure 2, arrow b). Likewise, the carbon cost of acquiring water (E_cost_) was predicted to increase with increasing drought severity (Figure 2, arrow c), as water becomes more limited and energetically expensive to extract under low soil moisture. We expected that the carbon cost of nitrogen acquisition would correlate positively with the carbon cost of acquiring nitrogen relative to water at the whole-plant scale (β_plant_) (Figure 2, arrow d), while the carbon cost of transpiration would correlate negatively with β_plant_ (Figure 2, arrow e). Furthermore, we propose that β_plant_ will at least partially predict the corresponding costs at the leaf level (β_leaf_) (Figure 2, arrow f) to predict χ (positive correlation, Figure 2, arrow g). Finally, we expected a negative correlation between χ and N_area_ (Figure 2, arrow h), and a positive correlation between N_area_ and V_cmax_ (Figure 2, arrow i), to maximize carbon assimilation at the lowest summed resource uptake and use cost.

**Figure 1.**
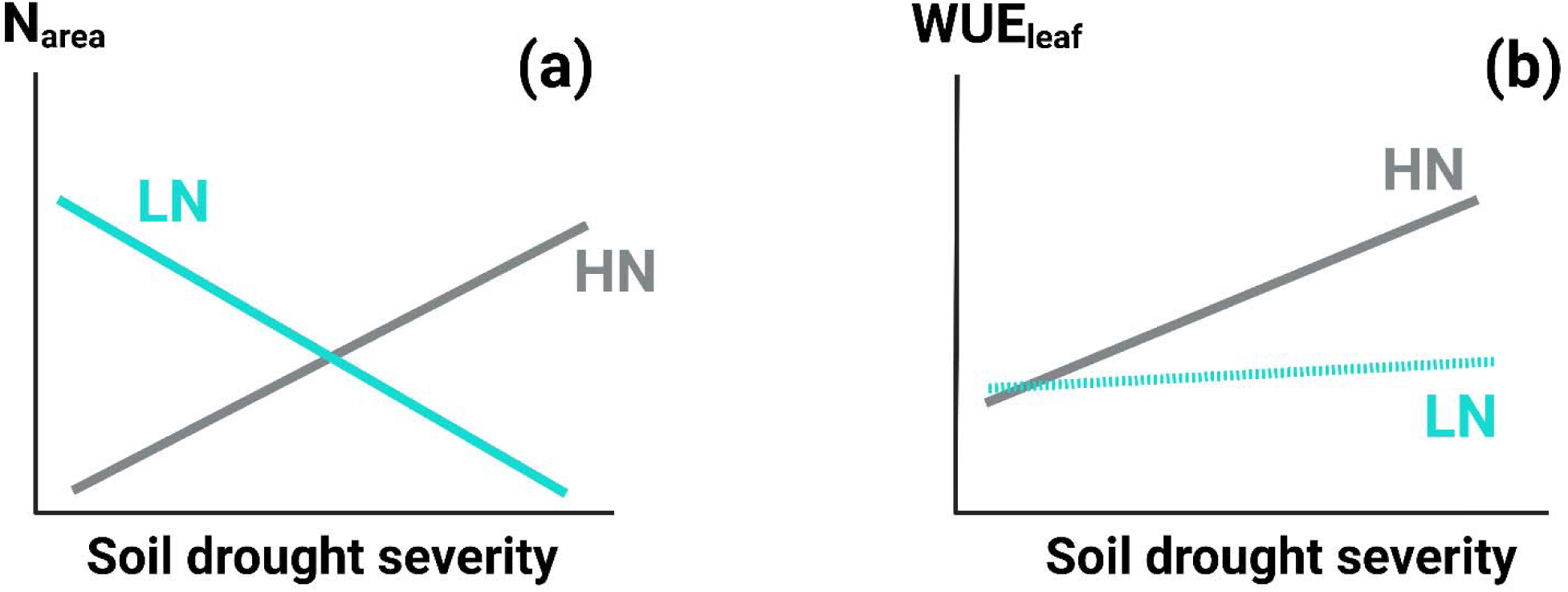
Hypothetical representation of the expected variation in leaf nitrogen on an area basis (N_area_) (a) and leaf water-use efficiency (WUE_leaf_) (b) with increasing soil drought severity. We expect that N_area_ will increase with soil drought severity to compensate for the partial closure of stomata and reduced intercellular CO_2_ concentration in plants grown under high nitrogen availability (HN), aligning with photosynthetic least-cost theory (a). However, plants grown under low nitrogen availability (LN) were expected to be unable to meet their leaf nitrogen demand, as reduced transpiration-driven mass flow, combined with low nitrogen concentrations in the soil, will hinder their nitrogen uptake—contrary to the expectations of photosynthetic least-cost theory. Thus, instead of increasing, their N_area_ is expected to decrease with soil drought severity (a). Consequently, because plants grown under HN were expected to align with the least-cost theory, their WUE_leaf_ was expected to be higher than that of plants grown under LN and to increase with soil drought severity (b). However, in plants grown under LN, WUE_leaf_ was not expected to vary with soil drought severity.

**Figure 2.**
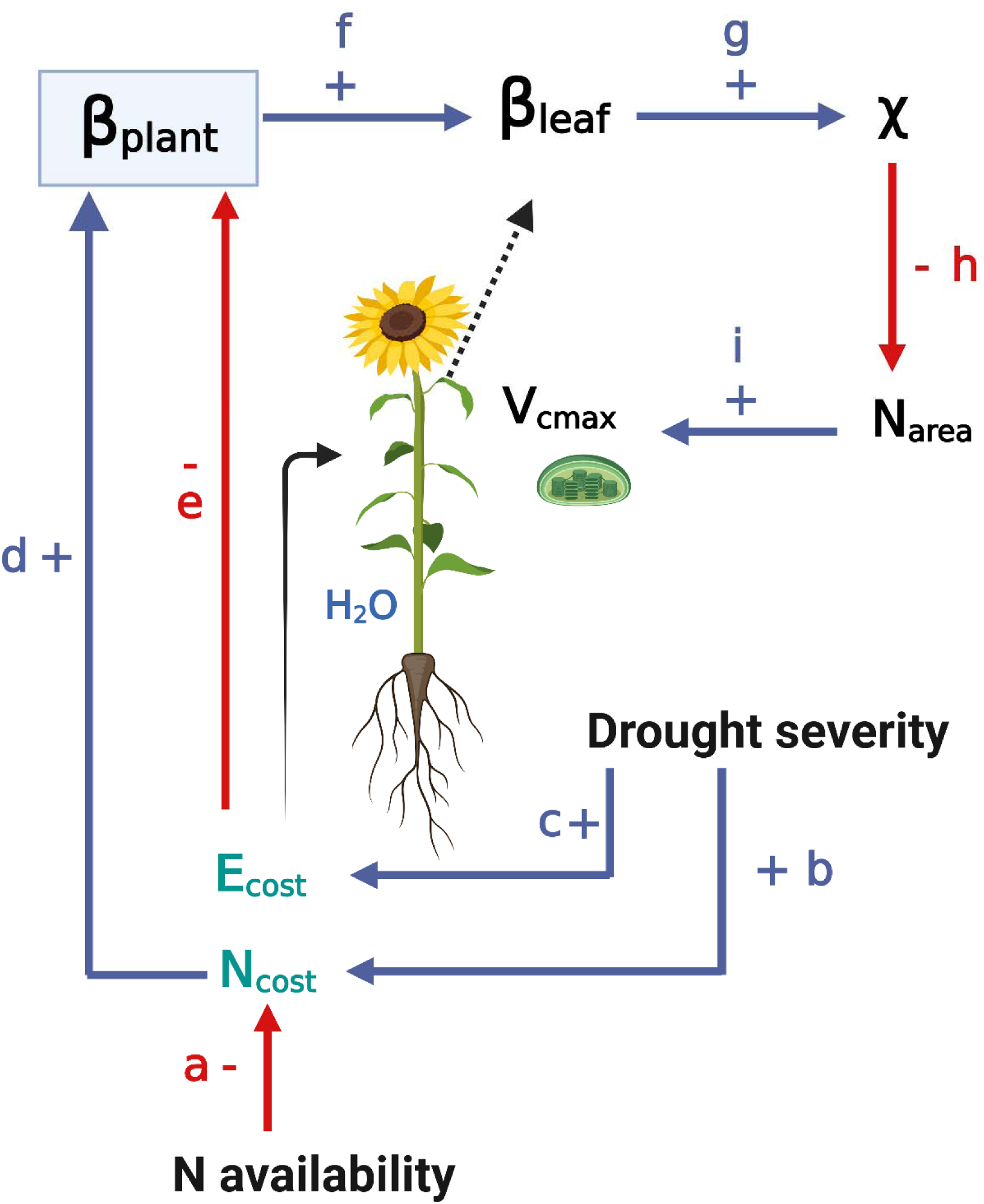
Hypothetical pathways linking the carbon costs of acquiring nitrogen relative to water at the whole-plant level (β_plant_) to those at the leaf level (β_leaf_) and their effects on leaf carbon economics. Increasing soil nitrogen availability is expected to decrease the carbon costs of nitrogen acquisition (N_cost_) (arrow a), while increasing soil drought severity, which reduces transpiration-driven mass flow, is expected to increase N_cost_ by hindering further nitrogen uptake (arrow b). Soil drought severity would also increase the cost of transpiration (E_cost_) (arrow c) as water becomes more costly to extract. Since the relative cost of nutrient acquisition to water acquisition (β_plant_) is defined as the ratio of N_cost_ to E_cost_, it is expected to increase with increasing N_cost_ (arrow d) and decreasing E_cost_ (arrow e). β_plant_ is expected to positively correlate with the relative costs of nutrient acquisition to water acquisition at the leaf level (β_leaf_) (arrow f), which is derived from χ (arrow g). Finally, χ is expected to negatively correlate with N_area_ (arrow h) and N_area_ is expected to correlate positively with V_cmax_ (arrow i), as predicted by photosynthetic least-cost theory.

## Materials and Methods

### Nitrogen and drought treatments

Annual common sunflower (*Helianthus annuus* L.) seeds were planted in 48 large white pots (18 liters each, Model S-7914W, Uline, USA) filled with a potting mixture of 65% natural sand (Pavestone sand) and 35% sphagnum peat moss (Premier Horticulture Inc, Quakertown, PA, USA). Immediately after germination, we divided the pots into two equal groups, thinned the plants to one per pot, and applied two levels of nitrogen fertilization (24 pots per group, corresponding to the two nitrogen treatments). The low nitrogen (LN) treatment consisted of a fertilization dose of 70 mg N L^-1^, while the high nitrogen (HN) treatment received 210 mg N L^-1^. A modified Hoagland solution (Hoagland & Arnon, 1950) was used, with equivalent concentrations of other macro- and micronutrients across treatments (Table S1). Each pot received 500 mL of the solution twice per week.

Approximately 30 days after germination, we conducted the first round of leaf trait and gas exchange measurements (see “Leaf traits” and “Gas exchange measurements” sections).

Immediately following these measurements, plants in both LN and HN treatments were randomly divided into three groups (8 plants per group) and assigned to one of three water treatments: Well-Watered (WW), Slow Progressive Drought (SPD), or Fast Developing Drought (FDD). The descriptions of each watering treatment are as follows:

- Well-Watered (WW): Plants were maintained at 100% field capacity (FC). The FC of the potting mixture was determined following the protocol described in the supplementary information document (See Notes S1 – Supplementary Information document).
- Slow Progressive Drought (SPD): Water was withheld until the potting mixture reached 50–60% FC. This level was maintained for approximately 12–15 days, after which water was withheld again until the mixture reached 40% FC. This final level was maintained until the end of the experiment. Differences in plant size and transpiration rates between HN and LN treatments (Figures 3a and 3b) resulted in variability in the time required to reach and maintain 50–60% FC or 40% FC across plants and nitrogen treatments (see “Drought Severity Index” section below).
- Fast Developing Drought (FDD): Water was withheld until the potting mixture reached 33% FC. This level was maintained for approximately 10–12 days before water was withheld again to reduce FC to 20%. This level was sustained until one week before the experiment’s conclusion. At that point, plants were re-wetted to 80% FC (Figures 3c and 3d). After re-wetting, water was again withheld to reduce FC to 20%. However, some plants, particularly those in the LN treatment, never reached this level by the experiment’s end (Figure 3d).

**Figure 3.**
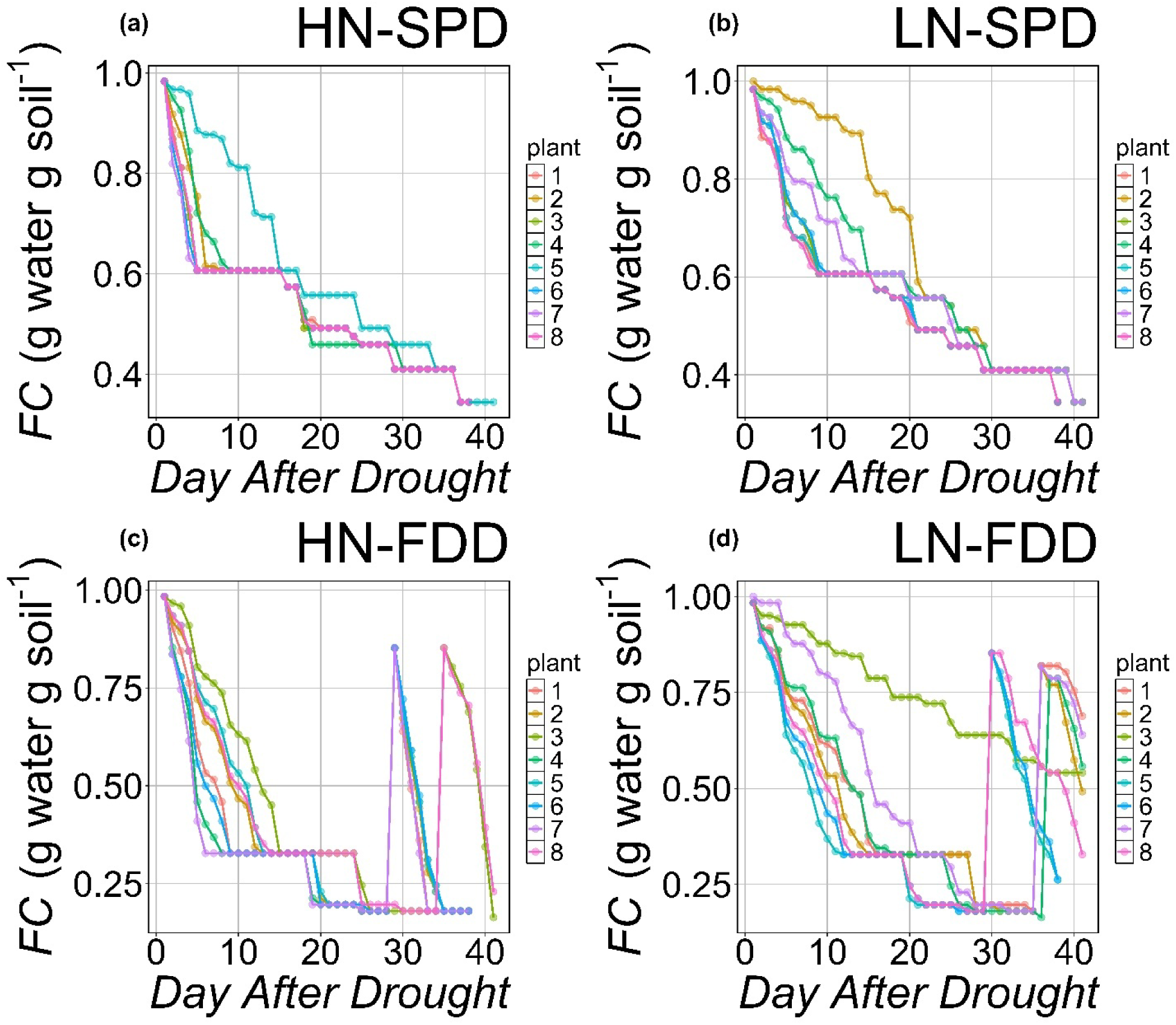
Progression of field capacity over time following drought treatments for plants under (a) high nitrogen level (HN) and slow progressive drought (SPD), (b) low nitrogen level (LN) and slow progressive drought (SPD), (c) high nitrogen level (HN) and fast-developing drought (FDD), and (d) low nitrogen level (LN) and fast-developing drought (FDD).

### Drought Severity Index (DSI)

Since plants varied in size depending on nitrogen fertilization levels, they also differed in transpiration rates. As a result, not all plants reached the target FC at the same time, leading to a lag between nitrogen treatments and even between plants within the same nitrogen treatment

(Figure 3). This lag was particularly evident in plants under LN (Figures 3b and 3d). To account for the individual drought history of each plant, we quantified drought impacts relative to each plant’s unique experience of drought progression. This was achieved by calculating a daily drought severity index (DSI_daily_) as the difference between the maximum soil field capacity (FC_max_) and the actual field capacity (FC) for a given day.:

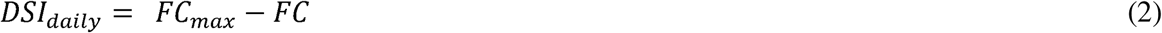

To account for the cumulative effect of drought overtime, we calculated Drought Severity Index (DSI) as the sum of daily drought severity index (DSI_daily_) values from the start of the drought application (t0) to each subsequent day (t), normalized by the number of days after drought application (DAD):

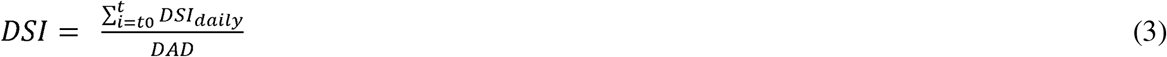

DSI is expressed in % day^-1^, reflecting the intensity and progression of soil moisture deficit over time.

### Cumulative nitrogen level calculation

For each plant, the cumulative nitrogen input (N level) was calculated daily by summing all nitrogen applications received from the beginning of the experiment up to the day of each measurement:

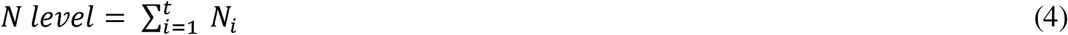

where N_i_ is the nitrogen applied on day *i*, and *t* is the day of each measurement. This approach provides a daily record of the total nitrogen supplied to each plant. N_i_ was calculated as the product of nitrogen levels in mg L^-1^ (70 for LN treatments and 210 for HN treatments) and the volume of solution added.

### Gas exchange measurements

Gas exchange measurements, leaf traits, and chlorophyll content were assessed at four time points: (1) before drought application (30 days after germination) and at three time points after drought application, (2) 17 days after drought application (DAD), (3) 30 DAD, and (4) 38 DAD. However, some plants, particularly those in the LN treatment, did not reach the target drought level at these times. For these plants that did not reach the target drought level, measurements were taken at 19–24 DAD, 35 DAD, and 41 DAD.

Gas exchange measurements were conducted using a Li-COR LI-6800 portable photosynthesis system equipped with a 6800-01A fluorometer head and a 6 cm² aperture (Li-COR Biosciences, Lincoln, NE, USA). For each plant, a young, fully expanded leaf was clamped into the LI-6800 cuvette to generate a CO_2_ response curve using the Dynamic Assimilation™ Technique (Saathoff & Welles, 2021). Reference CO_2_ concentrations ranged from 20 to 1620 μmol mol ¹, with a ramp rate of 200 μmol mol ¹ min ¹. Data were logged every 5 seconds, generating 96 points per response curve. The cuvette environment was maintained under saturated light (2000 μmol m ² s ¹), constant temperature (25°C), and a leaf vapor pressure deficit (VPD) of 1.5 kPa. Leaves were allowed to acclimate until steady-state stomatal conductance and net photosynthesis (A_net_) were reached at 420 μmol mol ¹ CO_2_ before generating curves.

The response curves were used to fit A_net_ as a function of intercellular CO_2_ (C) using the “fitaci” function in the R package plantecophys (Duursma, 2015). From these fits, the maximum rate of RuBisCO carboxylation (V_cmax_, μmol CO_2_ m ² s ¹) and the maximum rate of electron transport for RuBP regeneration (J_max_, μmol CO_2_ m ² s ¹) were estimated based on the Farquhar, Von Caemmerer & Berry (1980) biochemical model of C_3_ photosynthesis. Triose phosphate utilization (TPU) limitations were included for most curves where A_net_ leveled out or declined at high CO_2_ concentrations. Snapshot A_net_ measurements were extracted from each A_net_/C_i_ curves at a CO_2_ concentration of 420 µmol mol^-1^ CO_2_ (A_net_; μmol CO_2_ m ² s ¹).

### Leaf traits

For each time-point, the focal leaves used for gas exchange measurements were punched to obtain eight 0.6 cm^2^ leaf disks using a cork borer. Four disks were selected for chlorophyll analysis and four others were selected to determine the leaf mass per area (LMA; g m^-2^). These two sets of fresh disks were scanned separately using a flatbed scanner, and their total areas were analyzed with ImageJ (Schneider *et al*., 2012) and the LeafArea R package (Katabuchi, 2015).

Disks selected for chlorophyll analysis were stored at -80°C, while the other four disks were dried in an oven at 65°C for at least 72 hours and weighed once a constant weight was achieved. LMA was calculated by dividing the dry biomass of the disks (g) by their fresh area (m^2^).

The dried disks from each individual were finely ground using a Mini-Beadbeater-24 (Biospec Products) and submitted to the Stable Isotope Facility (SIF) at the University of California, Davis, CA. Analyses of ^13^C and ^15^N discrimination were conducted using an Elementar Vario EL Cube (Elementar Analysensysteme GmbH, Langenselbold, Germany) coupled to a continuous-flow isotope ratio mass spectrometer (PDZ Europa 20-20 IRMS, Sercon Ltd, Cheshire, UK). Leaf nitrogen content on a mass basis (N_mass_; gN g^-1^) and carbon isotope discrimination (δ¹³C_leaf_; ‰) were obtained.

Leaf nitrogen content per area (N_area_; gN m^-2^) was calculated as:

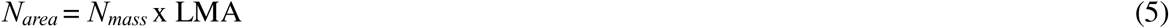

Leaf water-use efficiency (WUE_leaf_; μmol CO_2_ μmol H_2_O^-1^) was calculated as the ratio of A_net_ to g_s_, and leaf nitrogen-use efficiency (NUE_leaf_; μmol CO_2_ g N^-1^ s^-1^) was calculated as the ratio of Anet to Narea.

We calculated the ratio of intracellular to extracellular CO_2_ (χ; Pa Pa^-1^) from δ^13^C_leaf_ following Farquhar *et al* (1989), and then calculated β_leaf_ from χ following Prentice *et al*. (2014) (See Notes S2 – Supplementary Information document).

### Chlorophyll content and leaf nitrogen allocation

Chlorophyll content was determined following the protocol described in the supplementary information document (See Notes S3 – Supplementary Information document).

Leaf nitrogen allocation to different photosynthetic and structural pools at the leaf scale was divided into four categories: (1) nitrogen allocated to the carboxylation enzyme RuBisCO (ρ_rubisco_, gN gN^-1^); (2) nitrogen allocated to proteins involved in bioenergetics, including Cyt *b/f* related to electron transport and photophosphorylation (ρ_bioenergetics_, gN gN^-1^), (3) nitrogen allocated to light-harvesting proteins (ρ_lightharvesting_, gN gN^-1^), and (4) nitrogen allocated to cell wall proteins and structure pools (ρ_structure_, gN gN^-1^). We calculated these different fractions following the methods described in Niinemets & Tenhunen (1997), Niinemets *et al* (1998), and Onoda *et al* (2017) as:

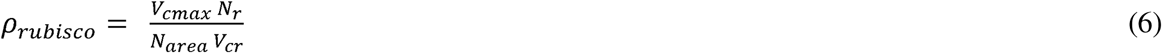

where Nr represents the amount of nitrogen in RuBisCO, assumed to be 0.16 gN g RuBisCO^-1^, and V_cr_ is the specific activity of Rubisco, assumed to be 20.5 µmol CO_2_ g RuBisCO ^−1^ s^−1^ at 25 °C.

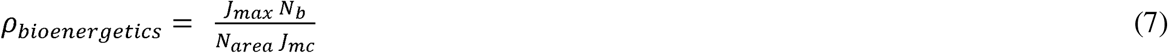

where N_b_ represents the amount of nitrogen in cytochrome f, assumed to be 0.1240695 gN µmol cytochrome f^-1^, and J_mc_ is the capacity of electron transport per cytochrome f unit, assumed to be 156 µmol electron µmol cytochrome f ^-1^ s^−1^

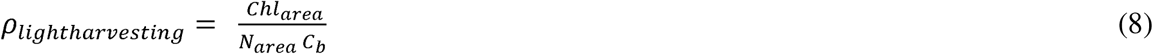

where Chl_area_ is the total chlorophyll content per unit area (mmol m^-2^), and Cb is the chlorophyll binding of the thylakoid protein complexes, assumed to be 2.75 mmol chlorophyll g chlorophyll N^-1^.

Finally, the proportion of leaf nitrogen content allocated to structural tissue (ρ_structure_; gN gN^−1^) was estimated as:

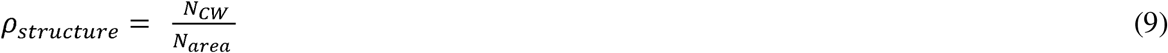

where N_cw_ is the leaf nitrogen content allocated to cell walls (gN m^−2^), calculated as a function of LMA using an empirical equation from Onoda *et al* (2017):

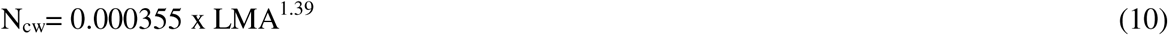

### Whole-plant measurements

Daily transpiration for each plant (E_plant_; g water day^-1^) was estimated by weighing pots and calculating the difference in weight between day *n* and day *n-1*. To minimize soil evaporation and ensure the measurement accounted solely for plant transpiration, the surface of each pot was covered with plastic film. We assumed that the plant biomass was negligible compared to the mass of the pot.

At the end of the experiment, all individuals were harvested immediately after the final gas exchange measurements. The fresh leaf area of all harvested leaves was measured using a flat-bed scanner and analyzed with ImageJ, as described for leaf disks in earlier measurements. Total fresh leaf area (cm^2^) was calculated as the sum of all individual leaf areas, including leaf disks used for assessing leaf traits and chlorophyll content.

The biomass of major organs (leaves, stems, roots, and flowers) was separated, oven-dried at 65°C for at least 72 hours, and weighed after reaching a constant weight. Total biomass (g) was calculated as the sum of the dry weights of leaves, stems, roots, and flowers.

Whole-plant carbon costs of nitrogen acquisition (N_cost_; g root g N leaf^-1^) were quantified as the ratio of root biomass to total leaf nitrogen:

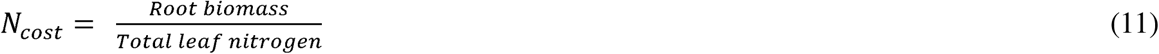

We estimated N_cost_ by assuming that additional unaccounted carbon allocation to root exudates, root respiration, fine root turnover, and mycorrhizal symbiosis scales proportionally to root biomass. Total leaf nitrogen was calculated as the product of N_mass_ and leaf biomass.

Similarly, whole-plant carbon costs for water acquisition (E_cost_; g root g water^-1^) were calculated as the ratio of root biomass to total water lost by each plant, which was estimated by summing daily transpiration over the entire experiment:

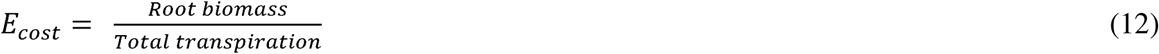

Finaly, we calculated the carbon cost of acquiring nitrogen relative to water (β_plant_) as the ratio of Ncost to Ecost (βplant=Ncost/Ecost).

### Data analysis

To test hypothesis 1, we analyzed leaf functional traits (χ, β_leaf_, LMA, N_mass_, and N_area_), and to test hypothesis 2 we analyzed leaf photosynthetic traits (V_cmax_, J_max_, A_net_, WUE_leaf_, NUEleaf, Chlarea), and leaf nitrogen allocation (ρrubisco, ρbioenergetics, ρlightharvesting, ρstructure) measured at the end of the experiment. We also calculated the rate of change in these traits with time and drought progression, relative to their pre-drought values. These traits were measured once before the drought (pre-drought) and at three subsequent time points during the drought period. The rate of change with time in each trait was calculated as:

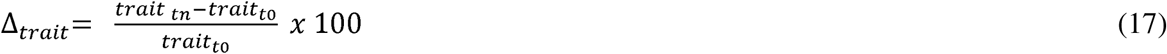

Where *t0* is the trait value at the predrought period, and *tn* is the trait value measured at time *n* after drought application.

Since plants experienced varying histories of drought progression and nitrogen accumulation, we analyzed these traits as functions of continuous variables - Drought Severity Index (DSI) and cumulative nitrogen (nitrogen level) - rather than categorical treatments (drought scenarios: WW, SPD, FDD; and nitrogen levels: HN vs. LN). Nonetheless, adopting drought severity progression effectively captured the drought scenarios, as plants in the SPD scenario accumulated less drought severity than those in the FDD scenario, as expected (Figure S1).

We conducted a series of linear mixed-effects models to examine how the drought severity index and cumulative nitrogen interacted to influence the rate of change of traits (Δ_trait_) over time. The models included drought severity index and cumulative nitrogen as well as their interaction as continuous fixed effects, while the identifier for each individual plant was included as a categorical random intercept term. We made sure that the climatic conditions (temperature and photosynthetically active radiation, PAR) were consistent across treatments in the greenhouse (Figure S2). Additionally, we applied the same models without the random intercept term to the final time-point measurements of leaf and photosynthetic traits, as well as whole-plant biomass and the total costs of nitrogen and water acquisition, to assess treatment effects on these variables.

We assessed residual normality for all models using Shapiro-Wilk tests of normality, comparing results for untransformed data with those using natural-log transformations (Shapiro-Wilk: p*>0.05* in all cases). Specifically, models for β_leaf_, N_mass_, N_area_, LMA, V_cmax_, J_max_, A_net_, Chl_area_, NUE_leaf_, WUEl_eaf_, plant surface area, and total biomass satisfied residual normality assumptions with log-transformed data, whereas other variables met these assumptions without transformation.

We fitted models without random terms using the “lm” function in R. For models with random effects, we used the ‘lmer’ function from the lme4 package (Bates et al., 2015). For all models, we calculated Type II Wald’s χ^2^ statistics to test the significance (a=0.05) of fixed effects using the Anova function in the ‘car’ package (Fox & Weisberg, 2019). Post-hoc comparisons were conducted using Tukey’s tests with the ‘emmeans’ package (Lenth *et al*., 2024), and degrees of freedom were approximated using the Kenward-Roger method. Linear regressions were plotted in all figures using estimated marginal means and standard errors across the range of DSI values for different cumulative nitrogen levels. All analyses and plots were performed in R version 4.1 (R Core Team, 2024).

Finally, to test hypothesis 3 (Figure 2), we performed a path analysis using a piecewise structural equation model. We constructed seven separate linear models: The first model predicted N_cost_ from DSI and nitrogen levels. The second model predicted E_cost_ from DSI. The third model predicted β_plant_from N_cost_ and E_cost_. The fourth model predicted β_leaf_from β_plant_, DSI and nitrogen levels. The fifth model predicted χ from β_leaf_. The sixth model predicted N_area_ from χ. The seventh model predicted V_cmax_ from N_area_. All models were built using the ‘lm’ function. These models were incorporated into the piecewise structural equation model using the ‘psem’ function in the ‘piecewiseSEM’ R package (Lifehack, 2016).

## Results

### Intracellular-to-ambient CO_2_ concentration ratio, leaf nitrogen, and leaf mass per area

As expected, 38–41 days after drought application, the intracellular-to-ambient CO_2_ concentration ratio (χ), and the relative costs of nutrient to water acquisition (β_leaf_), calculated from χ, decreased with increasing drought severity index (DSI) (*p < 0.05* in all cases; Table 1, Figures 4a and 4b). χ and β_leaf_ also decreased with increasing soil nitrogen levels (*p < 0.05* in both cases; Table 1, Figures 4a and 4b). A significant interaction between DSI and soil nitrogen levels revealed a greater decline in Δχ and Δβ_leaf_ with DSI as soil nitrogen levels increased (*p < 0.001* in both cases; Table S2 and Table S5, Figures S3a and S3b).

**Figure 4.**
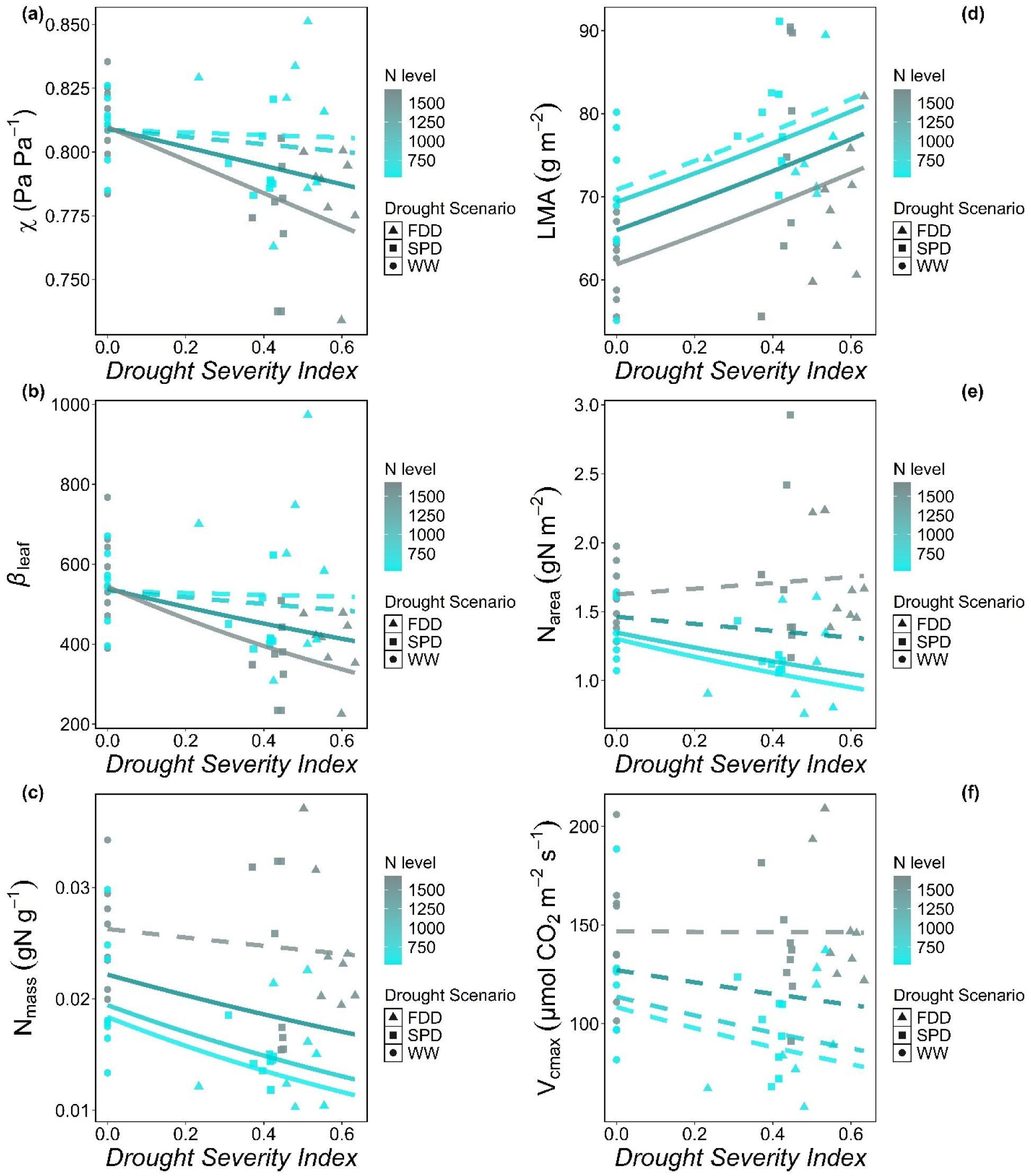
Fitted linear regression plots depicting the correlations between the drought severity index and (a) χ, (b) β_leaf_, (c) N_mass_, (d) LMA, (e) N_area_, and (f) V_cmax_ The regressions are derived from linear models for each dependent variable, with drought severity index and N level as interacting continuous effects. Circles represent well-watered plants, squares represent plants under the slow progressive drought scenario (SPD), and triangles represent plants under the fast-developing drought scenario (FDD). Separate lines are plotted for each N level, with color gradients ranging from cyan to grey indicating N levels from 650 mg per plant to 1680 mg N per plant. Solid lines indicate statistically significant trends (*p < 0.05*).

**Table 1.** ANOVA results for the linear models analyzing leaf trait variables measured at the end of the experiment after 38 to 41 days of drought application: χ (Pa Pa^1^), β_leaf_ (unitless), N_mass_ (gN g^1^), LMA (g m^2^), and N_area_ (gN m^2^) *.

| | df | $\chi$ | | $\beta_{\text{leaf}}$ | | $N_{\text{mass}}$ | | LMA | | $N_{\text{area}}$ | |
| --- | --- | --- | --- | --- | --- | --- | --- | --- | --- | --- | --- |
|  |  | F-value | P-value | F-value | P-value | F-value | P-value | F-value | P-value | F-value | P-value |
| Nitrogen level (N level) | 1 | 4.761 | **0.034** | 4.664 | **0.036** | 37.076 | *<0.001* | 8.575 | **0.005** | 29.078 | *<0.001* |
| Drought Severity Index (DSI) | 1 | 8.78 | **0.005** | 8.455 | **0.006** | 6.828 | **0.012** | 12.528 | *<0.001* | 1.137 | 0.286 |
| N level × DSI | 1 | 2.989 | 0.084 | 3.127 | 0.084 | 2.452 | 0.125 | 0.034 | 0.856 | 3.604 | 0.058 |

\*P-values < 0.001 are italicized and bolded, P-values < 0.05 are bolded. N level represents the cumulative nitrogen added to each plant from the beginning of fertilization treatments to the day of measurement (continuous. mg plant<sup>-1</sup>), Drought Severity Index (DSI) is the cumulative drought severity normalized by the number of days after drought application (continuous, % day<sup>-1</sup>), N level × DSI denotes interactions between nitrogen level and DSI. The sample size was 48.

As expected, 38–41 days after drought application, leaf nitrogen content on a mass basis (N_mass_) decreased with increasing DSI (*p < 0.05*; Table 1, Figure 4c). Increasing soil nitrogen levels also increased N_mass_ (*p < 0.001*; Table 1, Figure 4c). There were no interactive effects between DSI and soil nitrogen levels on N_mass_ (Table 1). Similarly, the percentage change over time in N_mass_ (ΔN_mass_) decreased with increasing DSI (*p < 0.05*; Table S2, Figure S3c).

Leaf mass per area (LMA) increased with increasing DSI (*p < 0.001*; Table 1, Figure 4d) and decreased with increasing soil nitrogen levels (*p < 0.05*; Table 1, Figure 4d). There were no interactive effects between DSI and nitrogen levels on LMA (Table 1). Similarly, the percentage change over time in LMA (ΔLMA) increased with increasing DSI (*p < 0.001*; Table S2), but with significant interactive effects with soil nitrogen levels. Post-hoc tests revealed a reduction in the positive relationship between LMA with DSI with decreasing soil nitrogen levels (Table S5, Figure S3d).

Leaf nitrogen content on an area basis (N_area_), calculated as the product of N_mass_ and LMA, increased with increasing soil nitrogen (*p < 0.001;* Table 1, Figure 4e). There was a marginally significant interaction between soil nitrogen levels and DSI (*p = 0.058*; Table 1) that indicated that DSI had an increasingly negative effect on N_area_ as soil nitrogen levels decreased (Figure 4e; Table 5), in line with hypothesis 1. The percentage change over time in N_area_ (ΔN_area_) increased with increasing DSI, with significant interactive effects with soil nitrogen levels (*p < 0.05*; Table S2) that indicated that the increase in ΔN_area_ with DSI increased with increasing soil nitrogen levels (Table S5, Figure S3e).

### Photosynthetic traits

Overall, after 38–41 days of drought application, V_cmax_ increased with increasing soil nitrogen levels (*p < 0.001*; Table 2, Figure 4f), but DSI did not affect significantly V_cmax_ (Table 2). In contrast, the percentage change in V_cmax_ over time (ΔV_cmax_) decreased with DSI, but this was lessened with increasing soil nitrogen levels (interaction: *p < 0.05*, Table S3, Table S5, Figure S3f). The observed patterns in J_max_ were similar to those observed for V_cmax_ (Tables 2, 5, S3, and S5).

**Table 2.**
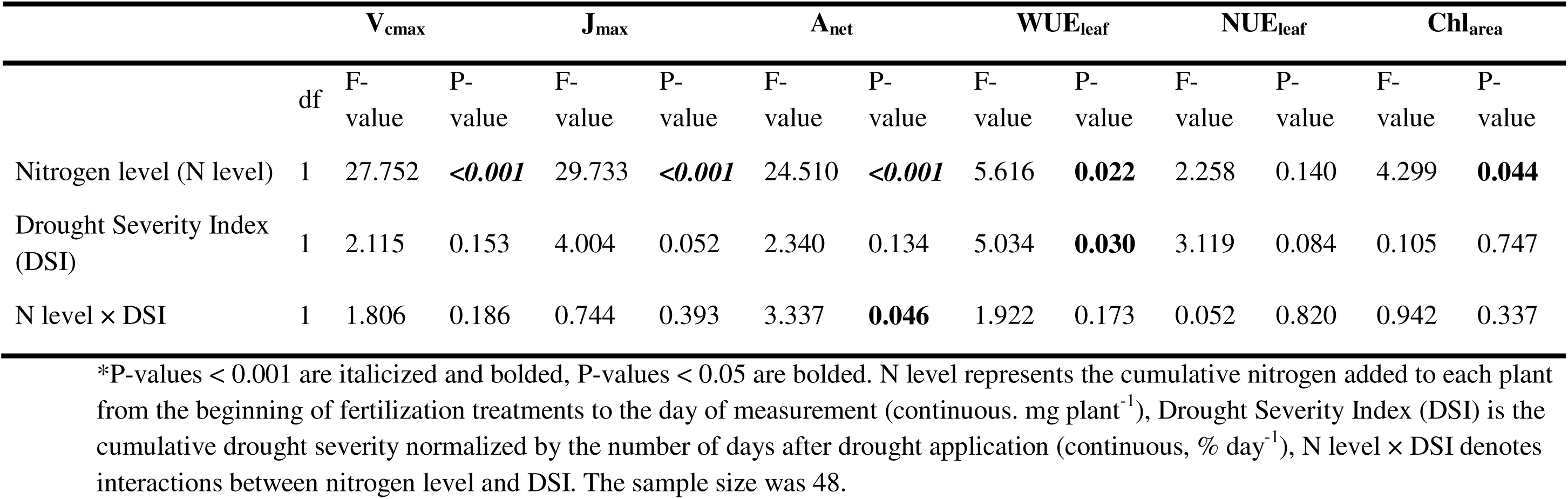
ANOVA results for the linear models analyzing photosynthetic trait variables measured at the end of the experiment after 38 to 41 days of drought application: V_cmax_ (µmol CO_2_ m^−2^ s^−1^), J_max_ (µmol CO_2_ m^−2^ s^−1^), A_net_ (µmol CO_2_ m^−2^ s^−1^), WUE_leaf_ (µmol CO_2_ µmol H_2_O ^−1^), NUE_leaf_ (µmol CO_2_ g N^−1^ s^−1^), and Chl_area_ (mmol m^−2^)*.

| | | $V_{\text{cmax}}$ | | $J_{\text{max}}$ | | $A_{\text{net}}$ | | $\text{WUE}_{\text{leaf}}$ | | $\text{NUE}_{\text{leaf}}$ | | $\text{Chl}_{\text{area}}$ | |
| --- | --- | --- | --- | --- | --- | --- | --- | --- | --- | --- | --- | --- | --- |
|  | df | F-value | P-value | F-value | P-value | F-value | P-value | F-value | P-value | F-value | P-value | F-value | P-value |
| Nitrogen level (N level) | 1 | 27.752 | ***<0.001*** | 29.733 | ***<0.001*** | 24.510 | ***<0.001*** | 5.616 | **0.022** | 2.258 | 0.140 | 4.299 | **0.044** |
| Drought Severity Index (DSI) | 1 | 2.115 | 0.153 | 4.004 | 0.052 | 2.340 | 0.134 | 5.034 | **0.030** | 3.119 | 0.084 | 0.105 | 0.747 |
| N level × DSI | 1 | 1.806 | 0.186 | 0.744 | 0.393 | 3.337 | **0.046** | 1.922 | 0.173 | 0.052 | 0.820 | 0.942 | 0.337 |

\*P-values $< 0.001$ are italicized and bolded, P-values $< 0.05$ are bolded. N level represents the cumulative nitrogen added to each plant from the beginning of fertilization treatments to the day of measurement (continuous. $\text{mg plant}^{-1}$ ), Drought Severity Index (DSI) is the cumulative drought severity normalized by the number of days after drought application (continuous, $\% \text{ day}^{-1}$ ), N level × DSI denotes interactions between nitrogen level and DSI. The sample size was 48.

After 38–41 days of drought application, the net photosynthetic rate (A_net_) increased with increasing soil nitrogen levels (*p < 0.001*; Table 2, Figure 5a), and decreased significantly with increasing DSI, but this effect was lessened as soil nitrogen levels increased (interaction: *p < 0.05*; Table 2, Figure 5a, Table 5). Similarly, the percentage change over time in A_net_ (ΔA_net_) significantly decreased with increasing DSI, but this effect was lessened by increasing soil nitrogen levels *(p < 0.05*, Table S3, Table S5, Figure S4a).

**Figure 5.**
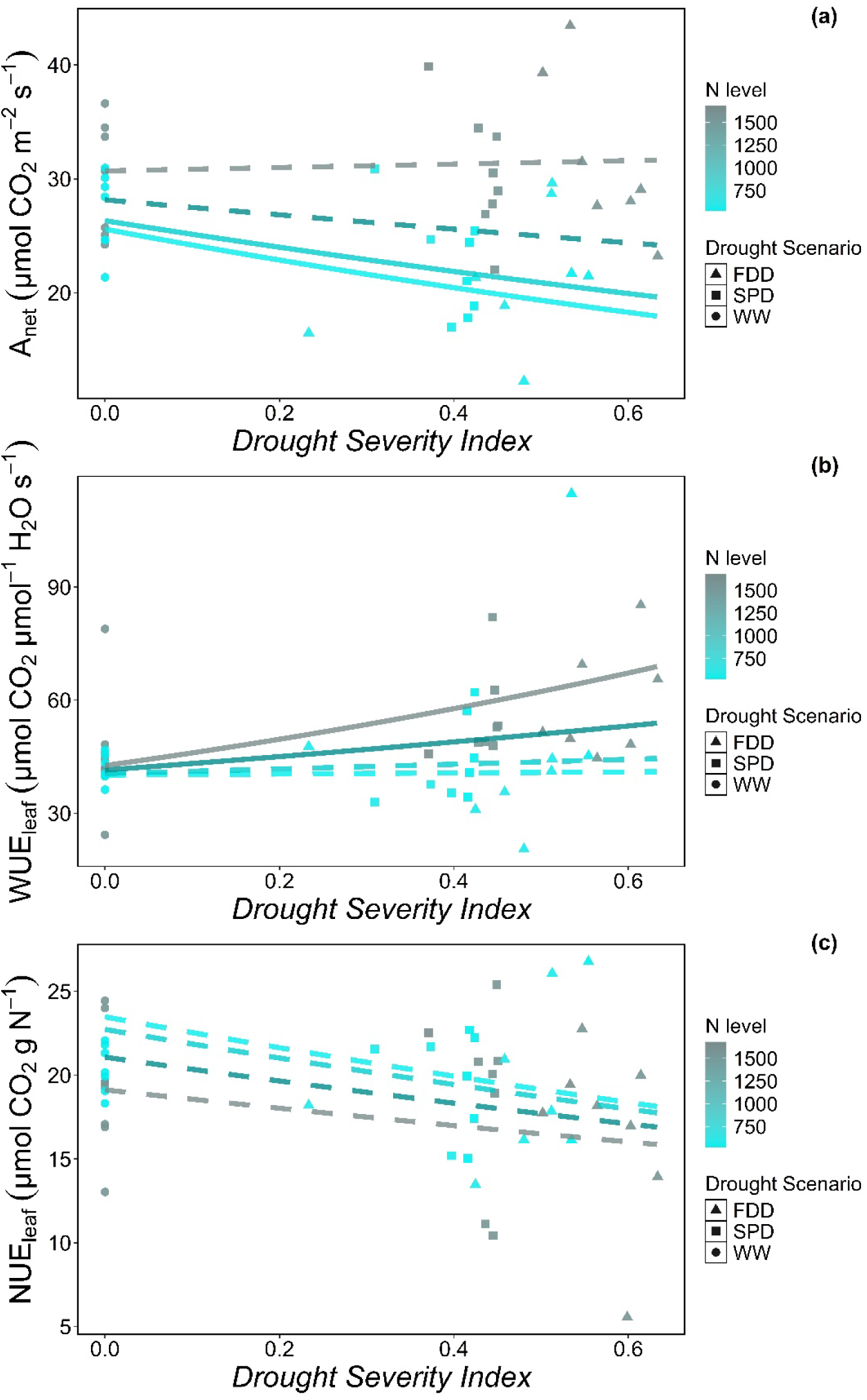
Fitted linear regression plots depicting the correlations between the drought severity index, (a) A_net_, (b) WUE_leaf_, and (c) NUE_leaf_. The regressions are derived from linear models for each dependent variable, with drought severity index and N level as interacting continuous effects. Circles represent well-watered plants, squares represent plants under the slow progressive drought scenario (SPD), and triangles represent plants under the fast-developing drought scenario (FDD). Separate lines are plotted for each N level, with color gradients ranging from cyan to grey indicating N levels from 650 mg per plant to 1680 mg N per plant. Solid lines indicate statistically significant trends (*p < 0.05*).

In line with hypothesis 2, leaf water-use efficiency (WUE_leaf_) after 38–41 days of drought application increased with increasing DSI and increasing soil nitrogen levels (*p < 0.05* for both cases; Table 2, Figure 5b). Similar patterns were observed in the percentage change over time in WUE_leaf_ (Tables S3 and S5, Figure S4b).

Leaf nitrogen use-efficiency (NUE_leaf_) was not significantly affected by soil nitrogen levels or DSI (Table 2, Figure 5c). However, the percentage change over time in NUE_leaf_ significantly increased with increasing soil nitrogen levels and decreased with increasing DSI (*p < 0.05* for both cases, Table S4, Figure S4c).

### Leaf chlorophyll content and leaf nitrogen partitioning

After 38–41 days of drought application, chlorophyll content per unit leaf area (Chl_area_) increased with increasing soil nitrogen levels (*p < 0.05*; Table 2). However, DSI had no significant effect on Chl_area_, nor was there an interaction between DSI and soil nitrogen levels (Table 2).

Nitrogen allocation to RuBisCO (ρ_rubisco_) and bioenergetics (ρ_bioenergetics_) was not significantly affected by either soil nitrogen levels or DSI (Table 3). However, their percentage change over time (Δρ_rubisco_ and Δρ_bioenergetics_) increased with increasing soil nitrogen levels (*p < 0.001* and *p < 0.05*, respectively; Table S4), but were unaffected by DSI (*p > 0.05*; Table S4). Nitrogen allocation to light-harvesting complexes (ρ_lightharvesting_) decreased with increasing soil nitrogen levels (*p < 0.001*; Table 3), but was unaffected by DSI (*p > 0.05*; Table 3).

**Table 3.**
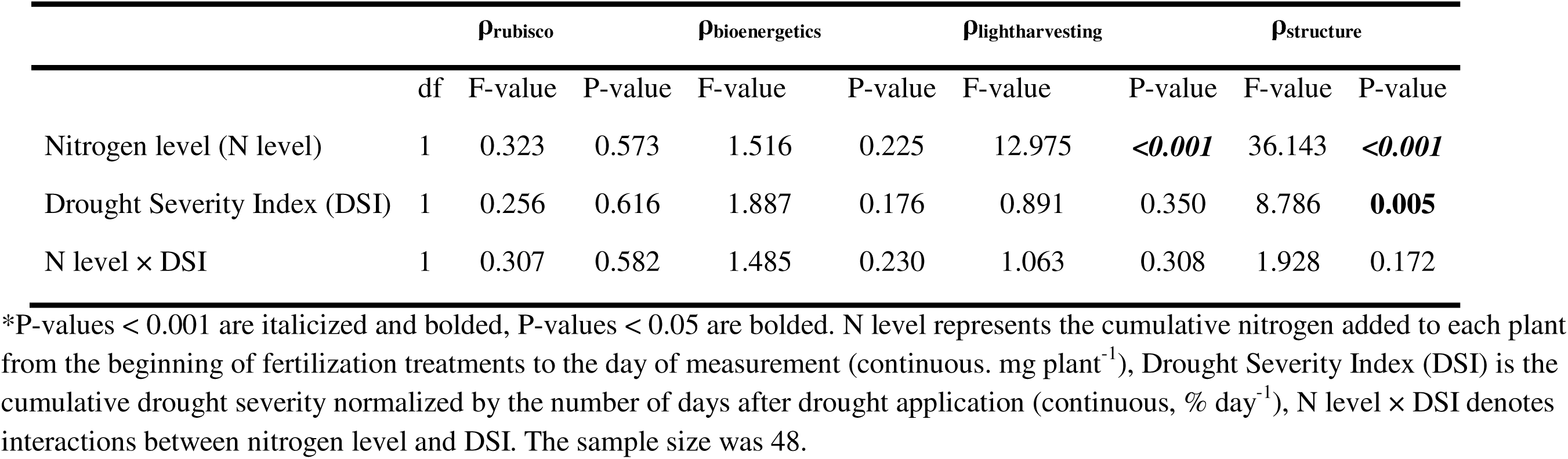
ANOVA results for the linear models analyzing leaf nitrogen allocation to different leaf pools at the end of the experiment after 38 to 41 days of drought application: ρ_rubisco_(gN gN^−1^), ρ_bioenergetics_ (gN g^1^), ρ_lightharvesting_ (gN gN 1), and ρ_structure_ (gN gN 1) *.

Nitrogen allocation to cell wall structure (ρ_structure_) decreased with increasing soil nitrogen levels (*p < 0.001*; Table 3) and increased with increasing DSI (*p < 0.05*; Table 3). The percentage change over time (Δρ_structure_) was only influenced by DSI (*p < 0.05*; Table S4), showing a significant increase with increasing DSI (Table S5).

### Whole-plant leaf area, biomass and resource acquisition costs

Plants receiving higher nitrogen levels had greater total leaf area (*p<0.001*; Table 4; Figures 6a). An interaction between DSI and nitrogen levels for total leaf area (*p<0.05*, Table 4) indicated a reduction in total leaf area with increasing DSI that increased in magnitude with increasing soil nitrogen levels (Figure 6a).

**Figure 6.**
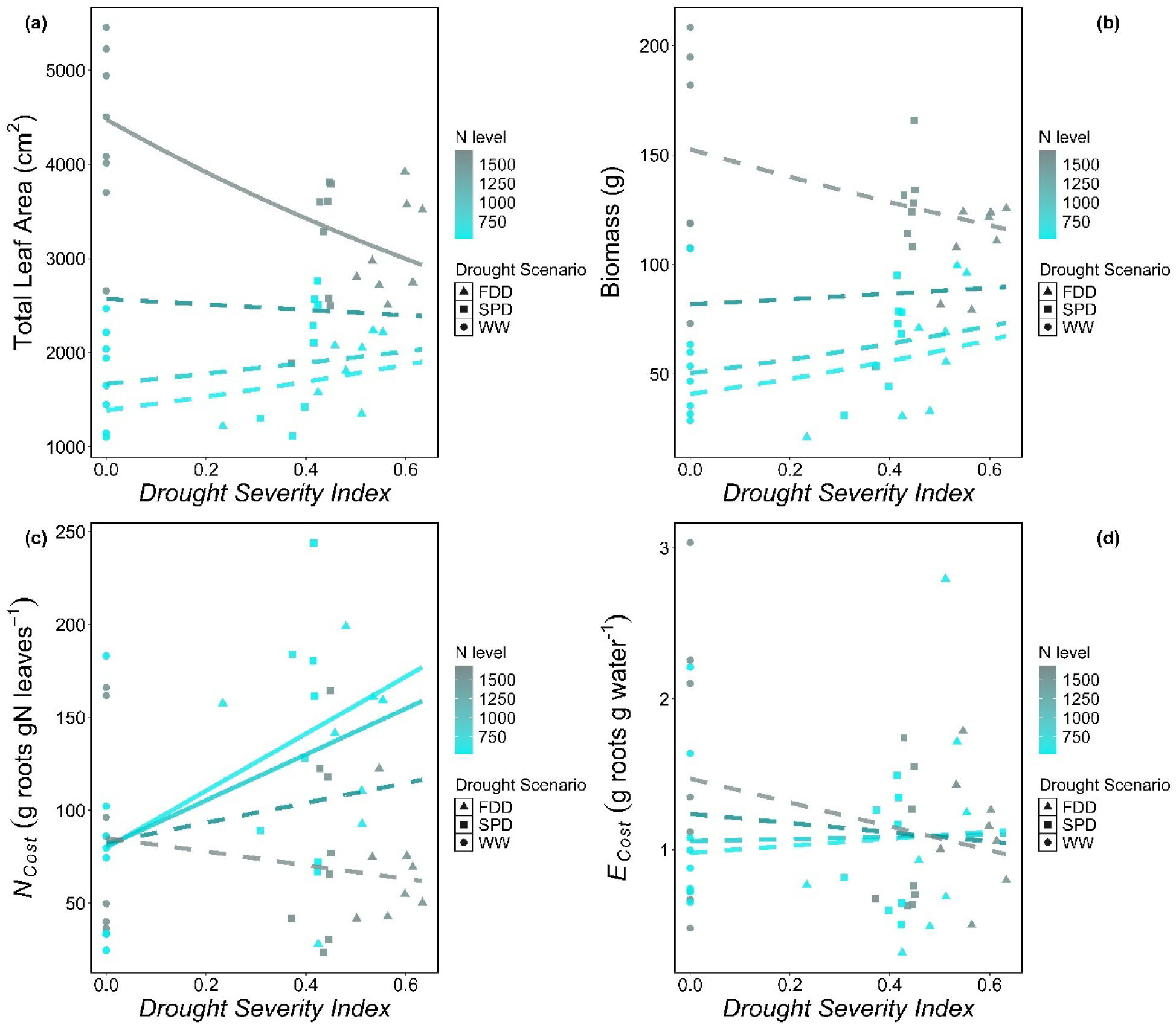
Fitted linear regression plots depicting the correlations between the drought severity index and (a) Total leaf Area, (b) Biomass, (c) E_cost_, and (d) N_cost_. The regressions are derived from linear models for each dependent variable, with drought severity index and N level as interacting continuous effects. Circle dots represent well-watered plants, square dots represent plants under the SPD scenario, and triangle dots represent plants under the FDD scenario. Separate lines are plotted for each N level, with color gradients ranging from cyan to grey indicating N levels from 650 mg per plant to 1680 mg N per plant. Solid lines indicate statistically significant trends (*p < 0.05*).

**Table 4.**
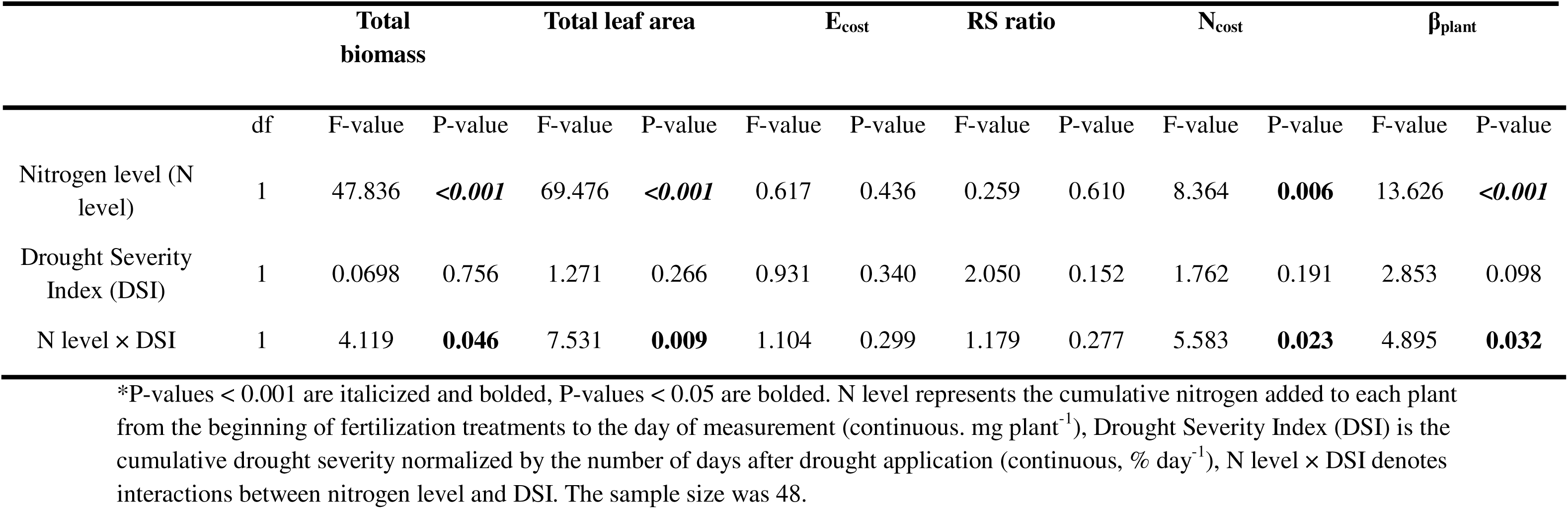
ANOVA results for the linear models analyzing whole-plant traits measured at the end of the experiment after 38 to 41 days of drought application: Total biomass (g), Total leaf area (cm^2^), E_cost_ (g roots g water^−1^), Root-to-shoot ratio (RS ratio), N_cost_ (g roots gN leaf^−1^), and β_plant_ (gN g water^-1^)*.

**Table 5.**
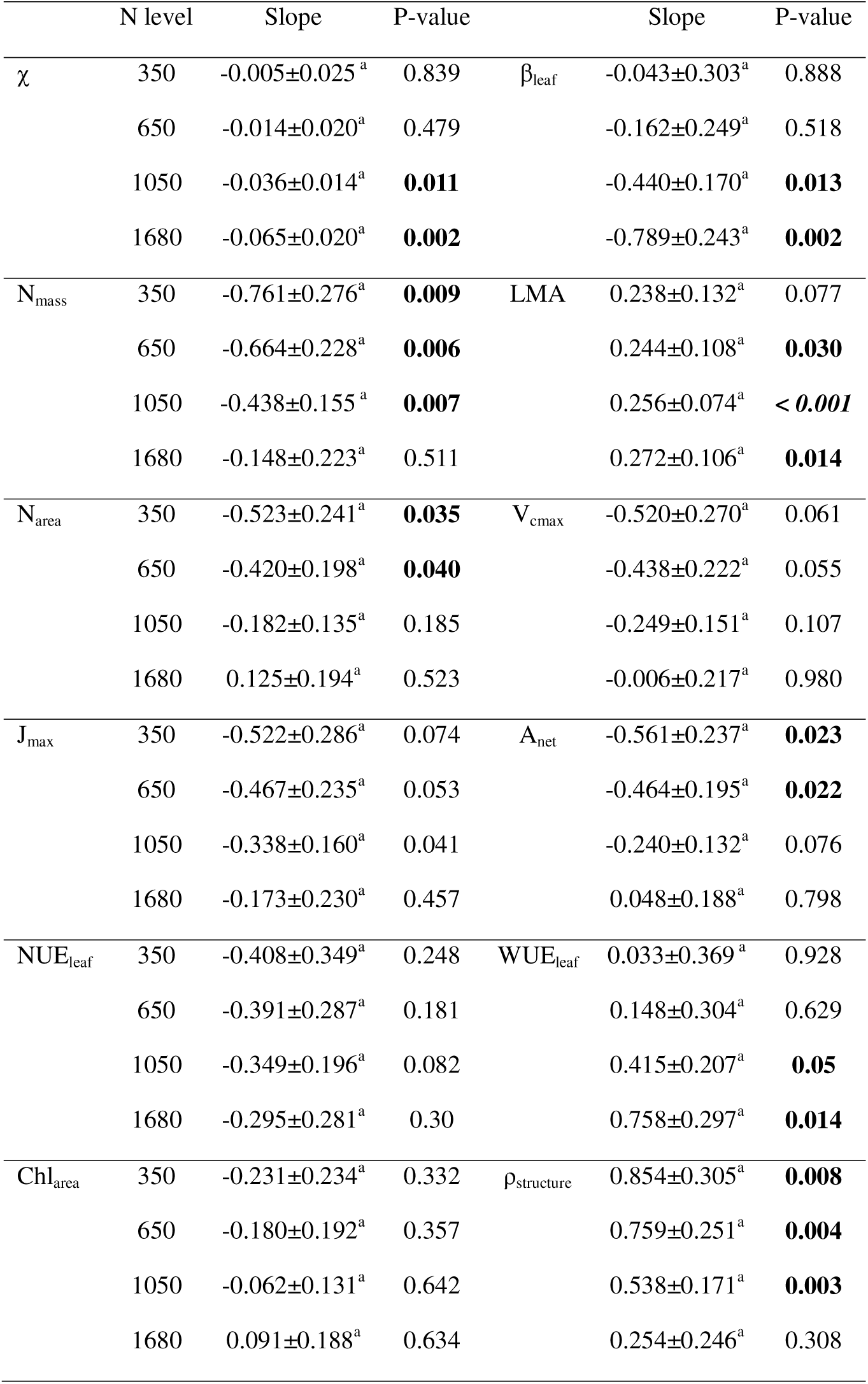
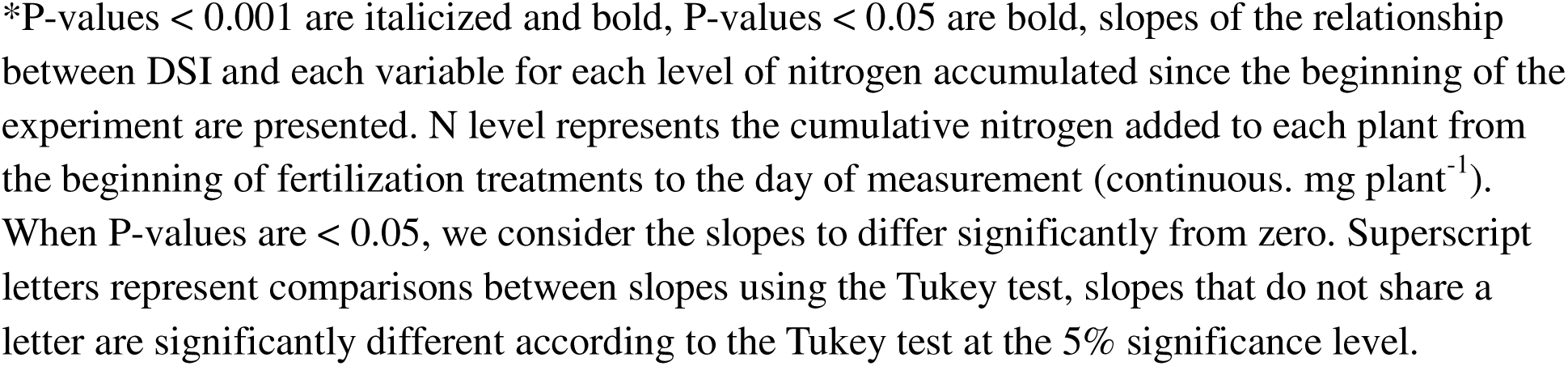
Post hoc analyses comparing the slopes of the relationships between drought severity index and χ (Pa Pa¹), βleaf (unitless), N_mass_ (gN g^1^), LMA (g m^2^), N_area_ (gN m^2^), V_cmax_ (µmol CO_2_ m^−2^ s^−1^), J_max_ (µmol CO_2_ m^−2^ s^−1^), A_net_ (µmol CO_2_ m^−2^ s^−1^), NUE_leaf_ (µmol CO_2_ g N^−1^ s^−1^), WUE_leaf_ (µmol CO_2_ µmol H_2_O ^−1^), Chl_area_ (mmol m^−2^), and ρ_structure_(gN gN 1), across soil nitrogen levels, calculated 38-41 days after drought application*.

Plants receiving higher nitrogen levels had greater biomass (*p<0.001*; Table 4; Figures 6b). There was a significant effect of the interaction between soil nitrogen levels and DSI on the total biomass (*p <0.05*, Table 4) that indicated that the positive relationship between soil nitrogen levels and biomass decreased with increasing DSI. There was no significant effect of DSI, soil nitrogen levels, or their interaction on the root-to-shoot ratio (Table 4).

As expected from hypothesis 3, N_cost_ increased with decreasing soil nitrogen levels (*p<0.05*: Table 4, Figure 6c). A significant interaction between soil nitrogen levels and DSI (*p<0.05*, Table 4) also revealed an increasingly positive relationship between N_cost_ and DSI as soil nitrogen levels decreased (Table 6, Figure 6c).

**Table 6.**
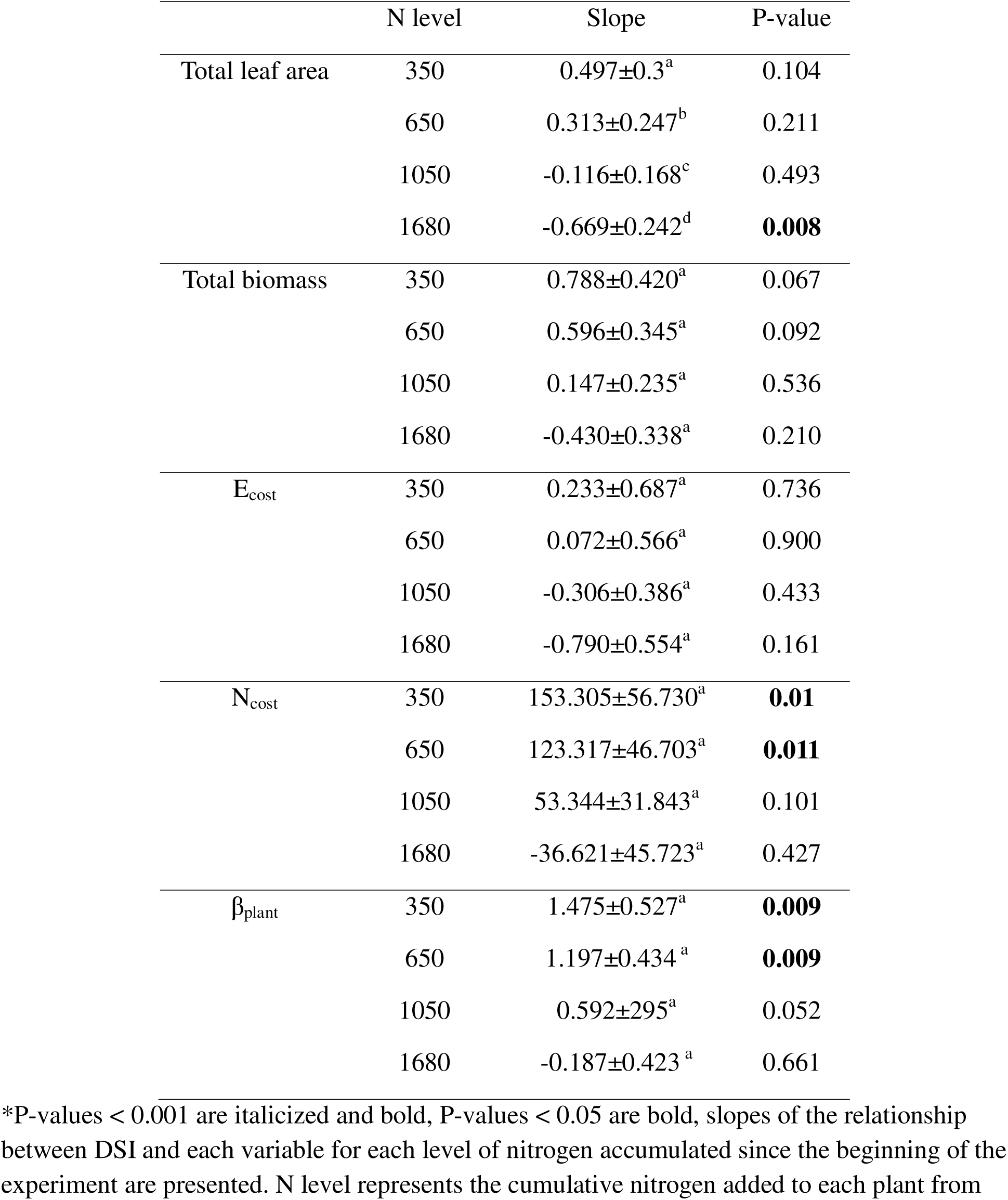
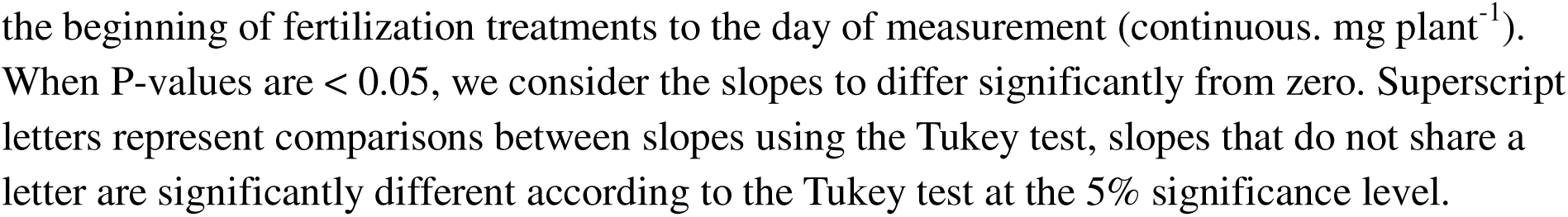
Post hoc analyses comparing the slopes of the relationships between drought severity index and the whole-plant total leaf area (cm^2^), total biomass (g), E_cost_ (g roots g water^−1^), N_cost_ (g roots gN leaf^−1^), and β_plant_ (gN g water^-1^) across soil nitrogen treatments, measured 38-41 days after drought application*.

|  | N level | Slope | P-value |
| --- | --- | --- | --- |
| Total leaf area | 350 | 0.497±0.3<sup>a</sup> | 0.104 |
|  | 650 | 0.313±0.247<sup>b</sup> | 0.211 |
|  | 1050 | -0.116±0.168<sup>c</sup> | 0.493 |
|  | 1680 | -0.669±0.242<sup>d</sup> | **0.008** |
| Total biomass | 350 | 0.788±0.420<sup>a</sup> | 0.067 |
|  | 650 | 0.596±0.345<sup>a</sup> | 0.092 |
|  | 1050 | 0.147±0.235<sup>a</sup> | 0.536 |
|  | 1680 | -0.430±0.338<sup>a</sup> | 0.210 |
| E<sub>cost</sub> | 350 | 0.233±0.687<sup>a</sup> | 0.736 |
|  | 650 | 0.072±0.566<sup>a</sup> | 0.900 |
|  | 1050 | -0.306±0.386<sup>a</sup> | 0.433 |
|  | 1680 | -0.790±0.554<sup>a</sup> | 0.161 |
| N<sub>cost</sub> | 350 | 153.305±56.730<sup>a</sup> | **0.01** |
|  | 650 | 123.317±46.703<sup>a</sup> | **0.011** |
|  | 1050 | 53.344±31.843<sup>a</sup> | 0.101 |
|  | 1680 | -36.621±45.723<sup>a</sup> | 0.427 |
| β<sub>plant</sub> | 350 | 1.475±0.527<sup>a</sup> | **0.009** |
|  | 650 | 1.197±0.434<sup>a</sup> | **0.009** |
|  | 1050 | 0.592±0.295<sup>a</sup> | 0.052 |
|  | 1680 | -0.187±0.423<sup>a</sup> | 0.661 |

\*P-values < 0.001 are italicized and bold, P-values < 0.05 are bold, slopes of the relationship between DSI and each variable for each level of nitrogen accumulated since the beginning of the experiment are presented. N level represents the cumulative nitrogen added to each plant from
the beginning of fertilization treatments to the day of measurement (continuous. mg plant<sup>-1</sup>). When P-values are < 0.05, we consider the slopes to differ significantly from zero. Superscript letters represent comparisons between slopes using the Tukey test, slopes that do not share a letter are significantly different according to the Tukey test at the 5% significance level.

Unexpectedly, DSI did not affect the carbon costs of water acquisition (E_cost_), and there were no significant effects of soil nitrogen levels or their interaction with DSI on E_cost_ (Table 4, Figure 6d). However, the whole-plant carbon costs of acquiring nitrogen relative to water (β_plant_) increased with decreasing soil nitrogen levels (*p < 0.001*; Table 4). A significant interaction between soil nitrogen levels and DSI (*p < 0.05*; Table 4) revealed that the positive relationship between β_plant_ and DSI increased with decreasing soil nitrogen levels (Table 6).

### Effects of whole-plant carbon costs on leaf economics

The path analysis (Figure 7) revealed that N_cost_ decreased with increasing soil nitrogen levels (standardized estimate coefficient = -0.38). However, contrary to our expectations, it was not significantly affected by DSI. Similarly, E_cost_ was not significantly affected by DSI. Unsurprisingly, β_plant_ was positively affected by N_cost_ (standardized estimate coefficient = 1.05) and negatively affected by E_cost_ (standardized estimate coefficient = -0.5). In line with our expectations, β_plant_ correlated positively with β_leaf_(standardized estimate coefficient = 0.33), and β_leaf_correlated positively with χ (standardized estimate coefficient = 0.96). As predicted by photosynthetic least-cost theory, χ correlated negatively with N_area_ (standardized estimate coefficient = -0.5), and N_area_ correlated positively with V_cmax_ (standardized estimate coefficient = 0.68). Finally, we did not find an effect of DSI on β_plant_. However, DSI had a direct negative effect on β_leaf (_standardized estimate coefficient = -0.42)

**Figure 7.**
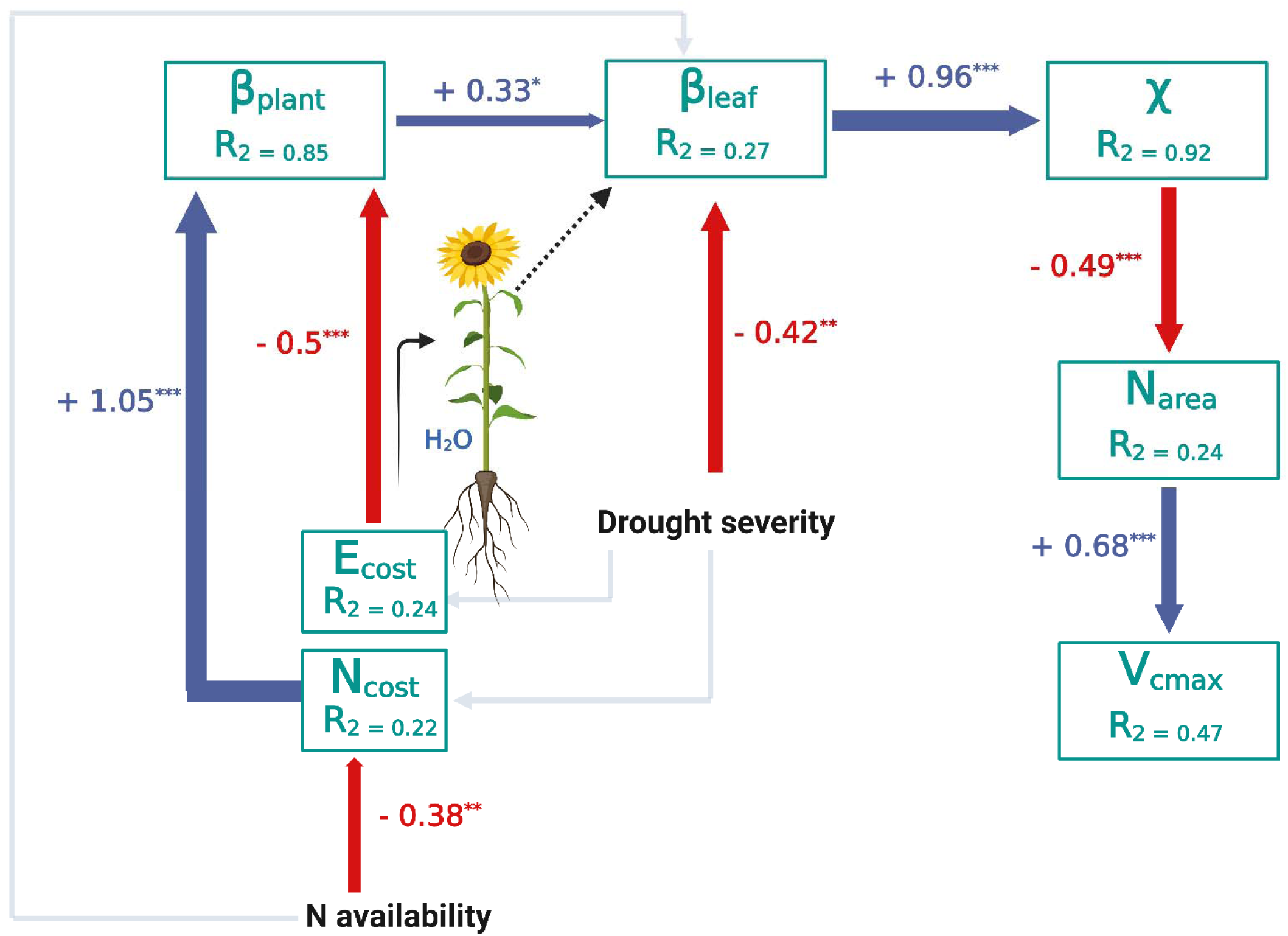
Structural equation model exploring how whole-plant carbon costs of acquiring nitrogen relative to water affect leaf-level economics. Arrows represent unidirectional relationships between variables. Path coefficients are depicted as simple standardized regression coefficients. The width of connections is based on estimates of standardized path coefficients, with solid lines denoting significant connections and semi-transparent lines indicating non-significant connections. Red arrows denote negative relationships, and dark blue arrows negatives ones. The coefficients of determination R^2^ for component models are given in the boxes of response variables.

## Discussion

Unable to move in search of water and nutrients, plants must adopt strategies and trade-offs to meet their needs (Rockwell & Sage, 2022). Photosynthetic least-cost theory proposes that plants trade nitrogen for water to maximize CO_2_ assimilation while minimizing the unit costs of nitrogen and water acquisition. By partially closing their stomata under water stress to conserve water—thereby increasing leaf water-use efficiency—plants upregulate RuBisCO content, leading to an increase in leaf nitrogen content and a subsequent decrease in leaf nitrogen-use efficiency. While this hypothesis has been supported at both regional (Prentice *et al*., 2014; Dong *et al*., 2020) and global scales (Wang *et al*., 2017) across species, less attention has been given to the interplay and interdependence between plant water relations and nutrient status within the same species. In this study, we explored this interplay by addressing two key questions: (1) How do soil water stress interaction with soil nitrogen limitation together influence this trade-off at the leaf level? (2) How do the whole-plant carbon costs of acquiring limiting resources shape leaf water and nitrogen economy?

### The least-cost theory does not hold under low soil water and low soil nitrogen availability

We found that the decline in leaf nitrogen on a mass basis (N_mass_) with increasing DSI was more pronounced under low soil nitrogen levels (Figure 4c). This suggests that rhizosphere depletion of dissolved nitrogen was not sufficiently compensated by nitrogen transport from the bulk soil via transpiration-driven mass flow to meet leaf nitrogen demand. Despite an increase in leaf mass per area (LMA) with soil drought severity across most soil nitrogen levels (Figure 4d), leaf nitrogen on an area basis (N_area_) decreased significantly under low soil nitrogen levels (Figure 4e) - contrary to the expectations of the least-cost theory, confirming hypothesis 1. In a recent study, Salazar Tortosa *et al*. (2018) proposed the theoretical framework of the “isohydric trap,” which suggests that transpiration-driven water flow is essential for nutrient uptake. Under drought conditions, this dependence can create a detrimental feedback loop in strictly isohydric species with tight stomatal control, leading to nutrient deficits and stoichiometric imbalances.

Their findings demonstrated that drought reduced nutrient uptake in mountain pine species with strict stomatal regulation of transpiration (i.e., strong isohydry), but not in anisohydric species. Our results align with these findings under low soil nitrogen levels, despite the fact that the common sunflower species used in our study is considered anisohydric, characterized by relaxed stomatal control and sustained transpiration (Tardieu & Simonneau, 1998). However, the traditional isohydric–anisohydric dichotomy has been revisited and refined, as evidence suggests that tight regulation of leaf water potential does not necessarily correlate with greater stomatal control in so-called isohydric species (Martínez Vilalta & Garcia Forner, 2017). Our results also align with other studies and meta-analyses (Fotelli *et al*., 2004; Hu *et al*., 2013; Joseph *et al*., 2021) that have reported drought-induced reductions in soil nitrogen uptake and plant nitrogen concentrations in deciduous tree species. Joseph *et al*. (2021) attributed this reduction primarily to a decrease in root surface area due to a loss of fine root biomass rather than to physiological limitations in nitrate and ammonium transporters. Additionally, they found that extreme drought reduced the allocation of new assimilates to roots, which they interpreted as a possible consequence of increased phloem sap viscosity, leading to reduced sieve tube conductivity (Dannoura *et al*., 2019).

While our experimental approach does not allow us to determine whether the decrease in leaf nitrogen content under drought and low soil nitrogen conditions is driven by reduced root surface area, fine root biomass loss, or declines in uptake and assimilation efficiency, we found that drought severity under low nitrogen conditions increased the carbon costs of nitrogen acquisition (Figure 6c). This suggests that, for the same carbon investment in root biomass, water-stressed plants took up less nitrogen than well-watered plants.

Under low nitrogen availability, the observed reduction in N_area_ with increasing soil drought severity correlated with a decline in V_cmax_ over time (Figure S3f), which in turn lowered A_net_ (Figure 5a). Consequently, this decrease in A_net_ mitigated the reduction in χ compared to plants grown under high nitrogen availability (Figure 4a). Since N_area_ decreased rather than increased with soil drought severity in plants grown under low nitrogen availability, WUE_leaf_ remained unchanged, in line with hypothesis 2. In contrast, under high soil nitrogen levels, the least-cost theory appears to hold, as N_area_, V_cmax_, and A_net_ remained similar in water-stressed plants relative to well-watered plants (Figure 4e and Figures 4f and 5a). Additionally, N_area_ even increased over time as drought progressed (Figure S3e). This increase, primarily driven by higher leaf mass per area (LMA, Figure 4d) rather than N_mass_, which showed a slight decline (Figure 4c), led to an overall increase in WUE_leaf_ with increasing DSI (Figure 5b), supporting hypothesis 2. These findings align with previous studies showing that nitrogen deposition increases WUE_leaf_ (Lu *et al*., 2019) and soil nitrogen addition enhances leaf nitrogen content and photosynthetic activity in grassland plants under extreme drought conditions, thereby improving drought resistance by delaying critical drought thresholds (Waraich *et al*., 2011; Shi *et al*., 2018; Song *et al*., 2019; He *et al*., 2024). Furthermore, our results are consistent with Ren *et al*. (2024), who reported that black locust leaves adapt to drought through increased LMA and moderate stomatal conductance reduction while maintaining constant N_area_ to optimize photosynthesis and carbon assimilation despite declining leaf N_mass_.

These findings highlight the importance of incorporating soil nitrogen availability into photosynthetic components of models that integrate photosynthetic least-cost optimality theory. When soil nitrogen availability is high, the least-cost hypothesis appears to be applicable.

However, under low nitrogen conditions, the direct effects of soil drought on plant nitrogen uptake must be explicitly considered and evaluated.

### Leaf nitrogen-use efficiency and nitrogen allocation to photosynthetic machinery are not affected by nitrogen levels or drought severity

Nitrogen partitioning between photosynthetic and non-photosynthetic proteins reflects a certain trade-off between carbon assimilation and environmental acclimation. Our findings indicate that neither leaf nitrogen-use efficiency (NUE_leaf_) nor the relative allocation of leaf nitrogen to key photosynthetic components (i.e., RuBisCO and bioenergetics, but see discussion of light harvesting below) was influenced by soil nitrogen levels or drought severity. These results are intriguing, as one might expect that increasing drought severity would enhance nitrogen allocation to RuBisCO and bioenergetics to compensate for reduced stomatal conductance and C_i_. Nonetheless, some studies have reported that soil water stress does not affect leaf nitrogen partitioning into photosynthetic pools or NUE_leaf_ (Zhong *et al*., 2019).

Additionally, previous research has shown that the proportion of leaf nitrogen allocated to photosynthesis decreases with increasing soil nitrogen availability (Waring *et al*., 2023) while NUE_leaf_ declines under high nitrogen availability (Zhong *et al*., 2019). However, these effects appear to be species-dependent, suggesting that sunflower leaf nitrogen allocation may not be significantly affected by the drought stress or soil nitrogen levels applied in our study.

Interestingly, we found that the proportion of nitrogen allocated to light harvesting significantly increased as soil nitrogen levels decreased. This aligns with the findings of Waring *et al*. (2023) but contrasts with Zhong *et al*. (2019) who reported that under low nitrogen availability, plants reduced nitrogen allocation to the light-harvesting system while increasing soluble protein and free amino acid levels or decreasing nitrogen allocation to the cell wall to maintain NUE_leaf_ under water deficit stress. In our study, however, plants grown under low nitrogen conditions increased nitrogen allocation to the cell wall. It is important to note that we did not measure nitrogen allocation to other leaf components, such as osmolytes and soluble proteins, which could further elucidate the observed differences.

### The carbon costs of nitrogen acquisition partially explained leaf-level economics

At the whole-plant level, as expected, total leaf area and biomass were consistently and positively influenced by soil nitrogen availability (Figures 6a and 6b). Nonetheless, the positive relationship between soil nitrogen levels and biomass decreased with increasing soil drought severity. Although drought severity reduced total leaf area in plants receiving the highest nitrogen levels, this did not significantly impact biomass. Instead, these plants compensated by producing leaves with higher LMA. Increasing LMA with drought severity is a well-documented response, allowing leaves to develop more rigid cell walls that enhance resistance to wilting (Poorter *et al*., 2009; Wellstein *et al*., 2017). However, the decline in A_net_ under drought stress and low nitrogen levels did not translate into biomass reductions. This may be attributed to increased respiration in well-watered plants compared to water-stressed ones. However, since we did not measure dark respiration at the leaf level or whole-plant respiration, a definitive interpretation remains unclear.

Supporting hypothesis 3, the whole-plant carbon costs of nitrogen acquisition (N_cost_) were higher in plants grown under low nitrogen availability compared to those receiving higher nitrogen levels (Figure 6c). These findings align with previous studies reporting increased carbon costs of nitrogen uptake under low soil nitrogen availability (Fisher *et al*., 2010; Shi *et al*., 2016; Perkowski *et al*., 2021). More importantly, N_cost_ increased with drought severity only under low nitrogen levels, primarily due to a decline in total leaf nitrogen rather than increased carbon allocation to roots, as root biomass remained unchanged under drought. This suggests that drought directly hinders nitrogen uptake and amplifies the carbon cost of nitrogen acquisition when nitrogen is scarce. We did not quantify total fine root biomass, root surface area, the flux of newly assimilated carbon to belowground tissues, or the physiological effects of drought on nitrate and ammonium transporter efficiency. Measuring these factors would have allowed us to disentangle the potential mechanisms contributing to reduced nitrogen uptake under drought and low soil nitrogen levels. However, path analysis across all treatments (all nitrogen levels combined) indicated that drought did not exert a direct effect on N_cost_ (Figure 7). This was likely because plants under high nitrogen levels, when analyzed separately, did not exhibit increased N_cost_ with increasing soil drought severity. Additionally, the path analysis revealed no effect of drought severity on the carbon costs of maintaining transpiration (E_cost_) (Figure 7), contrary to our expectations. This suggests that water-stressed plants did not allocate more carbon to root growth for water extraction, consistent with some studies indicating that root biomass does not always respond to drought as predicted by plant-optimality principles and that severe drought can even significantly hinder root growth (Eghball & Maranville, 1993; Wang *et al*., 2021).

At the whole-plant level, the relative costs of nutrient and water acquisition (β_plant_) correlated only partially with the corresponding leaf-level costs (β_leaf_) derived from χ. Interestingly, β_leaf_was directly affected by soil drought, but not via E_cost_, suggesting a decoupling between whole-plant E_cost_ and transpiration costs at the leaf level. This discrepancy could be interpreted in two ways: either the E_cost_ estimation at the whole-plant level is overly simplistic— being based solely on the root-to-transpiration ratio without accounting for sapwood respiration, fine root production, and other water transport costs—or the cost of transpiration at the leaf level is influenced by additional factors beyond water transport, such as the carbon costs of synthesizing osmoprotectants (Singh *et al*., 2015). Previous studies indicate that drought stress shifts the allocation of recent assimilates in favor of osmolyte pools, and plants exude relatively labeled carbon compounds into the rhizosphere to enhance water flow toward the roots (Hasibeder *et al*., 2015; Rabbi *et al*., 2018; Williams & De Vries, 2020; Wang *et al*., 2021)

Finally, the path analysis confirmed that χ correlated negatively with N_area_, while N_area_ correlated positively with V_cmax_ consistent with photosynthetic least-cost theory. To our knowledge, this is the first study linking whole-plant carbon costs of resource acquisition to estimates at the leaf level. Our findings emphasize the importance of both direct and indirect effects of soil resource availability in shaping leaf water and carbon economics under interacting drought and nitrogen conditions.

## Conclusion

Our findings showed that photosynthetic least-cost theory applies under high soil nitrogen availability but does not hold under severe drought and low soil nitrogen availability. Under low nitrogen, drought decreased N_area_, reducing A_net_ without increasing WUE_leaf_. In contrast, under high nitrogen, plants maintained N_area_, improving WUE_leaf_ in alignment with least-cost expectations. At the whole-plant level, drought increased the carbon costs of nitrogen acquisition under low nitrogen, highlighting a direct constraint of water limitation on nitrogen uptake. Our results emphasize the need to integrate the direct effects of drought severity on nitrogen uptake, especially under low soil nitrogen availability, into models simulating soil resource impacts on photosynthesis.

## References

Barber SA. 1962. A diffusion and mass-flow concept of soil nutrient availability. Soil Science 93: 39–49.

Barber SA. 1966. The role of root interception, mass-flow and diffusion in regulating the uptake of ions by plants from soil. In: Limiting steps in ion uptake by plants from soil.

Bista DR, Heckathorn SA, Jayawardena DM, Boldt JK. 2020. Effect of drought and carbon dioxide on nutrient uptake and levels of nutrient uptake proteins in roots of barley. American Journal of Botany 107: 1401–1409.

Bista DR, Heckathorn SA, Jayawardena DM, Mishra S, Boldt JK. 2018. Effects of drought on nutrient uptake and the levels of nutrient-uptake proteins in roots of drought-sensitive and- tolerant grasses. Plants 7: 28.

Brodribb T. 1996. Dynamics of changing intercellular CO_2_ concentration (ci) during drought and determination of minimum functional ci. Plant Physiology 111: 179–185.

Buckley TN. 2005. The control of stomata by water balance. New Phytologist 168: 275–292.

Buckley TN, Sack L, Farquhar GD. 2017. Optimal plant water economy. Plant, Cell & Environment 40: 881–896.

Cabrera-Bosquet L, Sánchez C, Araus JL. 2009. Oxygen isotope enrichment (Δ^18^ O) reflects yield potential and drought resistance in maize. Plant, Cell & Environment 32: 1487–1499.

Chapman N, Miller AJ, Lindsey K, Whalley WR. 2012. Roots, water, and nutrient acquisition: let’s get physical. Trends in Plant Science 17: 701–710.

Cheaib A, Waring EF, McNellis R, Perkowski EA, Martina JP, Seabloom EW, Borer ET, Wilfahrt PA, Dong N, Prentice IC. 2025. Soil nitrogen supply exerts largest influence on leaf nitrogen in environments with the greatest leaf nitrogen demand. Ecology letters 28: e70015.

Condon AG. 2004. Breeding for high water-use efficiency. Journal of Experimental Botany 55: 2447–2460.

Cowan IR. 1978. Stomatal behaviour and environment. In: Advances in botanical research. Elsevier, 117–228.

Cramer MD, Hawkins H-J, Verboom GA. 2009. The importance of nutritional regulation of plant water flux. Oecologia 161: 15–24.

Cramer MD, Hoffmann V, Verboom GA. 2008. Nutrient availability moderates transpiration in *Ehrharta calycina*. New Phytologist 179: 1048–1057.

Dannoura M, Epron D, Desalme D, Massonnet C, Tsuji S, Plain C, Priault P, Gérant D. 2019. The impact of prolonged drought on phloem anatomy and phloem transport in young beech trees (T Holtta, Ed.). Tree Physiology 39: 201–210.

Domingues TF, Meir P, Feldpausch TR, Saiz G, Veenendaal EM, Schrodt F, Bird M, Djagbletey G, Hien F, Compaore H, et al. 2010. Co limitation of photosynthetic capacity by nitrogen and phosphorus in West Africa woodlands. Plant, Cell & Environment 33: 959–980.

Dong N, Prentice IC, Wright IJ, Evans BJ, Togashi HF, Caddy-Retalic S, McInerney FA, Sparrow B, Leitch E, Lowe AJ. 2020. Components of leaf trait variation along environmental gradients. New Phytologist 228: 82–94.

Du B, Haensch R, Alfarraj S, Rennenberg H. 2024. Strategies of plants to overcome abiotic and biotic stresses. Biological Reviews 99: 1524–1536.

Duursma RA. 2015. Plantecophys - An R Package for analyzing and modelling leaf gas exchange data (PC Struik, Ed.). PLOS ONE 10: e0143346.

Edwards EJ, Osborne CP, Strömberg CAE, Smith SA, C Grasses Consortium, Bond WJ, Christin P-A, Cousins AB, Duvall MR, Fox DL, et al. 2010. The origins of C_4_ grasslands: integrating evolutionary and ecosystem science. Science 328: 587–591.

Eghball B, Maranville JW. 1993. Root development and nitrogen influx of corn genotypes grown under combined drought and nitrogen stresses. Agronomy Journal 85: 147–152.

Evans JR. 1989. Photosynthesis and nitrogen relationships in leaves of C3 plants. Oecologia 78: 9–19.

Evans JR, Clarke VC. 2019. The nitrogen cost of photosynthesis. Journal of Experimental Botany 70: 7–15.

Evans JR, Seemann JR. 1989. The allocation of protein nitrogen in the photosynthetic apparatus: costs, consequences, and control. Photosynthesis 8: 183–205.

Fan B, Westerband AC, Wright IJ, Gao P, Ding N, Ai D, Tian T, Zhao X, Sun K. 2024. Shifts in plant resource use strategies across climate and soil gradients in dryland steppe communities. Plant and Soil 497: 277–296.

Farquhar GD, Ehleringer JR, Hubic KT. 1989. Carbon isotope discrimination and photosynthesis. 40: 503–537.

Farquhar GD, Von Caemmerer S, Berry JA. 1980. A biochemical model of photosynthetic CO_2_ assimilation in leaves of C3 species. Planta 149: 78–90.

Ferreira Domingues T, Ishida FY, Feldpausch TR, Grace J, Meir P, Saiz G, Sene O, Schrodt F, Sonké B, Taedoumg H, et al. 2015. Biome-specific effects of nitrogen and phosphorus on the photosynthetic characteristics of trees at a forest-savanna boundary in Cameroon. Oecologia 178: 659–672.

Fisher JB, Sitch S, Malhi Y, Fisher RA, Huntingford C, Tan S -Y. 2010. Carbon cost of plant nitrogen acquisition: A mechanistic, globally applicable model of plant nitrogen uptake, retranslocation, and fixation. Global Biogeochemical Cycles 24: 2009GB003621.

Flexas J, Diaz-Espejo A, Gago J, Gallé A, Galmés J, Gulías J, Medrano H. 2014. Photosynthetic limitations in Mediterranean plants: A review. Environmental and Experimental Botany 103: 12–23.

Fotelli MN, Rienks M, Rennenberg H, Ge□ler A. 2004. Climate and forest management affect 15 N-uptake, N balance and biomass of European beech seedlings. Trees - Structure and Function 18: 157–166.

Franklin O, Harrison SP, Dewar R, Farrior CE, Brännström Å, Dieckmann U, Pietsch S, Falster D, Cramer W, Loreau M, et al. 2020. Organizing principles for vegetation dynamics. Nature Plants 6: 444–453.

Harrison SP, Cramer W, Franklin O, Prentice IC, Wang H, Brännström Å, De Boer H, Dieckmann U, Joshi J, Keenan TF, et al. 2021. Eco evolutionary optimality as a means to improve vegetation and land surface models. New Phytologist 231: 2125–2141.

Hasibeder R, Fuchslueger L, Richter A, Bahn M. 2015. Summer drought alters carbon allocation to roots and root respiration in mountain grassland. New Phytologist 205: 1117–1127.

He M, Dijkstra FA. 2014. Drought effect on plant nitrogen and phosphorus: a meta analysis. New Phytologist 204: 924–931.

He Y, Zhang R, Li P, Men L, Xu M, Wang J, Niu S, Tian D. 2024. Nitrogen enrichment delays the drought threshold responses of leaf photosynthesis in alpine grassland plants. Science of The Total Environment 913: 169560.

Hoagland DR, Arnon DI. 1950. The water culture method for growing plants without soil. Circular 347, 39 pp.

Hu B, Simon J, Kuster TM, Arend M, Siegwolf R, Rennenberg H. 2013. Nitrogen partitioning in oak leaves depends on species, provenance, climate conditions and soil type. Plant Biology 15: 198–209.

Joseph J, Luster J, Bottero A, Buser N, Baechli L, Sever K, Gessler A. 2021. Effects of drought on nitrogen uptake and carbon dynamics in trees (P Millard, Ed.). Tree Physiology 41: 927–943.

Katabuchi M. 2015. *LeafArea* : an R package for rapid digital image analysis of leaf area. Ecological Research 30: 1073–1077.

Kirnak H, Tas I, Kaya C, Higgs D. 2002. Effects of deficit irrigation on growth, yield and fruit quality of eggplant under semi-arid conditions. Australian journal of agricultural research 53: 1367–1373.

Kreuzwieser J, Gessler A. 2010. Global climate change and tree nutrition: influence of water availability. Tree Physiology 30: 1221–1234.

Lambers H, Oliveira RS. 2019. Mineral Nutrition. In: Plant Physiological Ecology. Cham: Springer International Publishing, 301–384.

Lambers H, Shane MW, Laliberté E, Swarts ND, Teste FP, Zemunik G. 2014. Plant mineral nutrition. Plant life on the sandplains in Southwest Australia, a global biodiversity hotspot: 101– 127.

Lawson T, Vialet-Chabrand S. 2019. Speedy stomata, photosynthesis and plant water use efficiency. New Phytologist 221: 93–98.

Leadley PW, Reynolds JF, Chapin Iii FS. 1997. A model of nitrogen uptake by Eriophorum vaginatum roots in the field: ecological implications. Ecological monographs 67: 1–22.

Lefcheck JS. 2016. PIECEWISESEM : Piecewise structural equation modelling in R for ecology, evolution, and systematics (R Freckleton, Ed.). Methods in Ecology and Evolution 7: 573–579.

Lenth RV, Bolker B, Buerkner P, Giné-Vázquez I, Herve M, Jung M, Love J, Miguez F, Riebl H, Singmann H. 2024. Package ‘emmeans’.

Lu X, Ju W, Jiang H, Zhang X, Liu J, Sherba J, Wang S. 2019. Effects of nitrogen deposition on water use efficiency of global terrestrial ecosystems simulated using the IBIS model. Ecological Indicators 101: 954–962.

Luo X, Keenan TF, Chen JM, Croft H, Colin Prentice I, Smith NG, Walker AP, Wang H, Wang R, Xu C, et al. 2021. Global variation in the fraction of leaf nitrogen allocated to photosynthesis. Nature Communications 12: 4866.

Martínez-Vilalta J, Garcia-Forner N. 2017. Water potential regulation, stomatal behaviour and hydraulic transport under drought: deconstructing the iso/anisohydric concept. Plant, Cell & Environment 40: 962–976.

Matimati I, Verboom GA, Cramer MD. 2014. Nitrogen regulation of transpiration controls mass-flow acquisition of nutrients. Journal of Experimental Botany 65: 159–168.

McMurtrie RE, Näsholm T. 2018. Quantifying the contribution of mass flow to nitrogen acquisition by an individual plant root. New Phytologist 218: 119–130.

Nielsen DR, Th. Van Genuchten M, Biggar JW. 1986. Water flow and solute transport processes in the unsaturated zone. Water resources research 22: 89S–108S.

Niinemets U, Kull O, Tenhunen JD. 1998. An analysis of light effects on foliar morphology, physiology, and light interception in temperate deciduous woody species of contrasting shade tolerance. Tree Physiology 18: 681–696.

Niinemets Ü, Tenhunen JD. 1997. A model separating leaf structural and physiological effects on carbon gain along light gradients for the shade tolerant species *Acer saccharum*. Plant, Cell & Environment 20: 845–866.

Nye PH, Marriott FHC. 1969. A theoretical study of the distribution of substances around roots resulting from simultaneous diffusion and mass flow. Plant and Soil 30: 459–472.

Onoda Y, Wright IJ, Evans JR, Hikosaka K, Kitajima K, Niinemets Ü, Poorter H, Tosens T, Westoby M. 2017. Physiological and structural tradeoffs underlying the leaf economics spectrum. New Phytologist 214: 1447–1463.

Paillassa J, Wright IJ, Prentice IC, Pepin S, Smith NG, Ethier G, Westerband AC, Lamarque LJ, Wang H, Cornwell WK, et al. 2020. When and where soil is important to modify the carbon and water economy of leaves. New Phytologist 228: 121–135.

Patterson TB, Guy RD, Dang QL. 1997. Whole-plant nitrogen- and water-relations traits, and their associated trade-offs, in adjacent muskeg and upland boreal spruce species. Oecologia 110: 160–168.

Peng Y, Bloomfield KJ, Cernusak LA, Domingues TF, Colin Prentice I. 2021. Global climate and nutrient controls of photosynthetic capacity. Communications Biology 4: 462.

Perkowski EA, Waring EF, Smith NG. 2021. Root mass carbon costs to acquire nitrogen are determined by nitrogen and light availability in two species with different nitrogen acquisition strategies (A Rogers, Ed.). Journal of Experimental Botany 72: 5766–5776.

Poorter H, Niinemets Ü, Poorter L, Wright IJ, Villar R. 2009. Causes and consequences of variation in leaf mass per area (LMA): a meta analysis. New Phytologist 182: 565–588.

Prentice IC, Dong N, Gleason SM, Maire V, Wright IJ. 2014. Balancing the costs of carbon gain and water transport: testing a new theoretical framework for plant functional ecology (J Penuelas, Ed.). Ecology Letters 17: 82–91.

Querejeta JI, Prieto I, Armas C, Casanoves F, Diémé JS, Diouf M, Yossi H, Kaya B, Pugnaire FI, Rusch GM. 2022. Higher leaf nitrogen content is linked to tighter stomatal regulation of transpiration and more efficient water use across dryland trees. New Phytologist 235: 1351–1364.

Rabbi SMF, Tighe MK, Flavel RJ, Kaiser BN, Guppy CN, Zhang X, Young IM. 2018. Plant roots redesign the rhizosphere to alter the three dimensional physical architecture and water dynamics. New Phytologist 219: 542–550.

Raschke K. 1976. How stomata resolve the dilemma of opposing priorities. Philosophical Transactions of the Royal Society of London. B, Biological Sciences 273: 551–560.

Reich PB. 2014. The world wide ‘fast–slow’ plant economics spectrum: a traits manifesto (H Cornelissen, Ed.). Journal of Ecology 102: 275–301.

Ren W, Tian L, Querejeta JI. 2024. Tight coupling between leaf Δ^13^ C and N content along leaf ageing in the N_2_ fixing legume tree black locust (*Robinia pseudoacacia* L.). Physiologia Plantarum 176: e14235.

Rengel Z. 1993. Mechanistic simulation models of nutrient uptake: a review. Plant and Soil 152: 161–173.

Rockwell F, Sage RF. 2022. Plants and water: the search for a comprehensive understanding. Annals of Botany 130: i–viii.

Saathoff AJ, Welles J. 2021. Gas exchange measurements in the unsteady state. Plant, Cell & Environment 44: 3509–3523.

Salazar-Tortosa D, Castro J, Villar-Salvador P, Viñegla B, Matías L, Michelsen A, Rubio De Casas R, Querejeta JI. 2018. The “isohydric trap”: A proposed feedback between water shortage, stomatal regulation, and nutrient acquisition drives differential growth and survival of European pines under climatic dryness. Global Change Biology 24: 4069–4083.

Sardans J, Peñuelas J. 2014. Hydraulic redistribution by plants and nutrient stoichiometry: Shifts under global change. Ecohydrology 7: 1–20.

Schneider CA, Rasband WS, Eliceiri KW. 2012. NIH Image to ImageJ: 25 years of image analysis. 9(7): 671–675.

Shi M, Fisher JB, Brzostek ER, Phillips RP. 2016. Carbon cost of plant nitrogen acquisition: global carbon cycle impact from an improved plant nitrogen cycle in the Community Land Model. Global Change Biology 22: 1299–1314.

Shi B, Wang Y, Meng B, Zhong S, Sun W. 2018. Effects of nitrogen addition on the drought susceptibility of the leymus chinensis meadow ecosystem vary with drought duration. Frontiers in Plant Science 9: 254.

Singh M, Kumar J, Singh S, Singh VP, Prasad SM. 2015. Roles of osmoprotectants in improving salinity and drought tolerance in plants: a review. Reviews in Environmental Science and Bio/Technology 14: 407–426.

Smith NG, Keenan TF. 2020. Mechanisms underlying leaf photosynthetic acclimation to warming and elevated CO_2_ as inferred from least cost optimality theory. Global Change Biology 26: 5202–5216.

Smith NG, Keenan TF, Colin Prentice I, Wang H, Wright IJ, Niinemets Ü, Crous KY, Domingues TF, Guerrieri R, Yoko Ishida F. 2019. Global photosynthetic capacity is optimized to the environment. Ecology letters 22: 506–517.

Song Y, Li J, Liu M, Meng Z, Liu K, Sui N. 2019. Nitrogen increases drought tolerance in maize seedlings. Functional Plant Biology 46: 350.

Stocker BD, Dong N, Perkowski EA, Schneider PD, Xu H, De Boer HJ, Rebel KT, Smith NG, Van Sundert K, Wang H, et al. 2025. Empirical evidence and theoretical understanding of ecosystem carbon and nitrogen cycle interactions. New Phytologist 245: 49–68.

Stocker BD, Wang H, Smith NG, Harrison SP, Keenan TF, Sandoval D, Davis T, Prentice IC. 2020. P-model v1. 0: An optimality-based light use efficiency model for simulating ecosystem gross primary production. Geoscientific Model Development 13: 1545–1581.

Tanguilig VC, Yambao EB, O’toole JC, De Datta SK. 1987. Water stress effects on leaf elongation, leaf water potential, transpiration, and nutrient uptake of rice, maize, and soybean. Plant and Soil 103: 155–168.

Tardieu F, Simonneau T. 1998. ariability among species of stomatal control under fluctuating soil water status and evaporative demand: modelling isohydric and anisohydric behaviours. : 419–432.

Torres-Ruiz JM, Cochard H, Delzon S, Boivin T, Burlett R, Cailleret M, Corso D, Delmas CEL, De Caceres M, Diaz-Espejo A, et al. 2024. Plant hydraulics at the heart of plant, crops and ecosystem functions in the face of climate change. New Phytologist 241: 984–999.

VanWallendael A, Soltani A, Emery NC, Peixoto MM, Olsen J, Lowry DB. 2019. A Molecular view of plant local adaptation: Incorporating stress-response networks. Annual Review of Plant Biology 70: 559–583.

Volaire F. 2018. A unified framework of plant adaptive strategies to drought: Crossing scales and disciplines. Global Change Biology 24: 2929–2938.

Volaire F, Gleason SM, Delzon S. 2020. What do you mean “functional” in ecology? Patterns versus processes. Ecology and Evolution 10: 11875–11885.

Voltas J, Romagosa I, Muñoz P, Araus JL. 1998. Mineral accumulation, carbon isotope discrimination and indirect selection for grain yield in two-rowed barley grown under semiarid conditions. European Journal of Agronomy 9: 147–155.

Wang R, Cavagnaro TR, Jiang Y, Keitel C, Dijkstra FA. 2021. Carbon allocation to the rhizosphere is affected by drought and nitrogen addition. Journal of Ecology 109: 3699–3709.

Wang H, Prentice IC, Keenan TF, Davis TW, Wright IJ, Cornwell WK, Evans BJ, Peng C. 2017. Towards a universal model for carbon dioxide uptake by plants. Nature Plants 3: 734–741.

Wankmüller FJP, Carminati A. 2023. Soil hydraulic constraints on stomatal regulation of plant gas exchange. In: Lüttge U, Cánovas FM, Risueño M-C, Leuschner C, Pretzsch H, eds. Progress in Botany. Progress in Botany Vol. 84. Cham: Springer Nature Switzerland, 327–350.

Waraich EA, Ahmad R, Ashraf MY, Saifullah, Ahmad M. 2011. Improving agricultural water use efficiency by nutrient management in crop plants. Acta Agriculturae Scandinavica, Section B-Soil & Plant Science 61: 291–304.

Waring EF, Perkowski EA, Smith NG. 2023. Soil nitrogen fertilization reduces relative leaf nitrogen allocation to photosynthesis (A Rogers, Ed.). Journal of Experimental Botany 74: 5166– 5180.

Wellstein C, Poschlod P, Gohlke A, Chelli S, Campetella G, Rosbakh S, Canullo R, Kreyling J, Jentsch A, Beierkuhnlein C. 2017. Effects of extreme drought on specific leaf area of grassland species: A meta analysis of experimental studies in temperate and sub Mediterranean systems. Global Change Biology 23: 2473–2481.

Westerband AC, Wright IJ, Maire V, Paillassa J, Prentice IC, Atkin OK, Bloomfield KJ, Cernusak LA, Dong N, Gleason SM, et al. 2023. Coordination of photosynthetic traits across soil and climate gradients. Global Change Biology 29: 856–873.

Westoby M, Falster DS, Moles AT, Vesk PA, Wright IJ. 2002. Plant ecological strategies: some leading dimensions of variation between species. Annual Review of Ecology and Systematics 33: 125–159.

Williams A, De Vries FT. 2020. Plant root exudation under drought: implications for ecosystem functioning. New Phytologist 225: 1899–1905.

Wright IJ, Reich PB, Westoby M. 2003. Least cost input mixtures of water and nitrogen for photosynthesis. The American Naturalist 161: 98–111.

Zhong C, Jian S-F, Huang J, Jin Q-Y, Cao X-C. 2019. Trade-off of within-leaf nitrogen allocation between photosynthetic nitrogen-use efficiency and water deficit stress acclimation in rice (Oryza sativa L.). Plant Physiology and Biochemistry 135: 41–50.

